# Cryo-EM structure of an active central apparatus

**DOI:** 10.1101/2022.01.23.477438

**Authors:** Long Han, Qinhui Rao, Renbin Yang, Yue Wang, Pengxin Chai, Yong Xiong, Kai Zhang

## Abstract

Accurately regulated ciliary beating in time and space is critical for diverse cellular activities, which impact the survival and development of nearly all eukaryotic species. An essential beating regulator is the conserved central apparatus (CA) of motile cilia, composed of a pair of microtubules (C1 and C2) associated with hundreds of protein subunits per repeating unit. It is largely unclear how CA plays its regulatory roles in ciliary motility. Here, we present high-resolution structures of *Chlamydomonas reinhardtii* CA by cryo-electron microscopy (cryo-EM) and its dynamic conformational behavior in multiple scales. The structures show how functionally related projection proteins of CA are clustered onto a spring-shaped scaffold of armadillo-repeat proteins, facilitated by elongated rachis-like proteins. The two halves of CA are brought together by elastic chain-like bridge proteins to achieve coordinated activities. We captured an array of kinesin-like protein (KLP1) in two different stepping states, which are actively correlated with beating wave propagation of cilia. These findings establish a structural framework for understanding the role of CA in cilia.

## Introduction

Motile cilia and flagella are evolutionarily conserved organelles responsible for the movement of individual cells and transport of extracellular materials through rhythmic beats (1). Ciliary motility is required for the survival of both unicellular organisms and high-level species (2–4). In humans, it plays essential roles in a variety of life activities, such as embryonic development, fertility, airway functions, and circulation of cerebral spinal fluid (5–9). Defects in ciliary structures and functions lead to numerous diseases termed ciliopathies, including congenital heart defects, hydrocephalus, and primary ciliary dyskinesia (PCD) (10–12).

The beating of cilia is driven by the axoneme, which is characterized by a ‘9+2’ scaffold structure containing nine peripheral microtubule doublets (MTDs) and a pair of central microtubules (C1 and C2) (13–15) (**Fig. 1a**). Each MTD binds two rows of minus-end-directed dynein motors, including inner-arm dyneins (IADs) and outer-arm dyneins (OADs), which generate the main mechanical forces required for ciliary beating via ATP hydrolysis (16–18). To achieve a rhythmic beat, ciliary dyneins are regulated by various axonemal components and extracellular signals (16, 19–21). The main regulator is the central apparatus (CA), composed of the two central microtubules and numerous associated proteins, including the plus-end-directed motor, kinesin-like protein (KLP1) (22, 23). Despite the existence of CA-less motile cilia in rare cases (24–27) or under extreme conditions (28), CA is essential for ciliary motility in those species that contain the CA structure (25, 29–31). It has been proposed that CA acts as a mechanical force distributor (32), which interacts with radial spokes to transmit mechanochemical signals to the nexin-dynein regulatory complexes and IADs for regulating OAD activities (33–38).

**Fig. 1:**
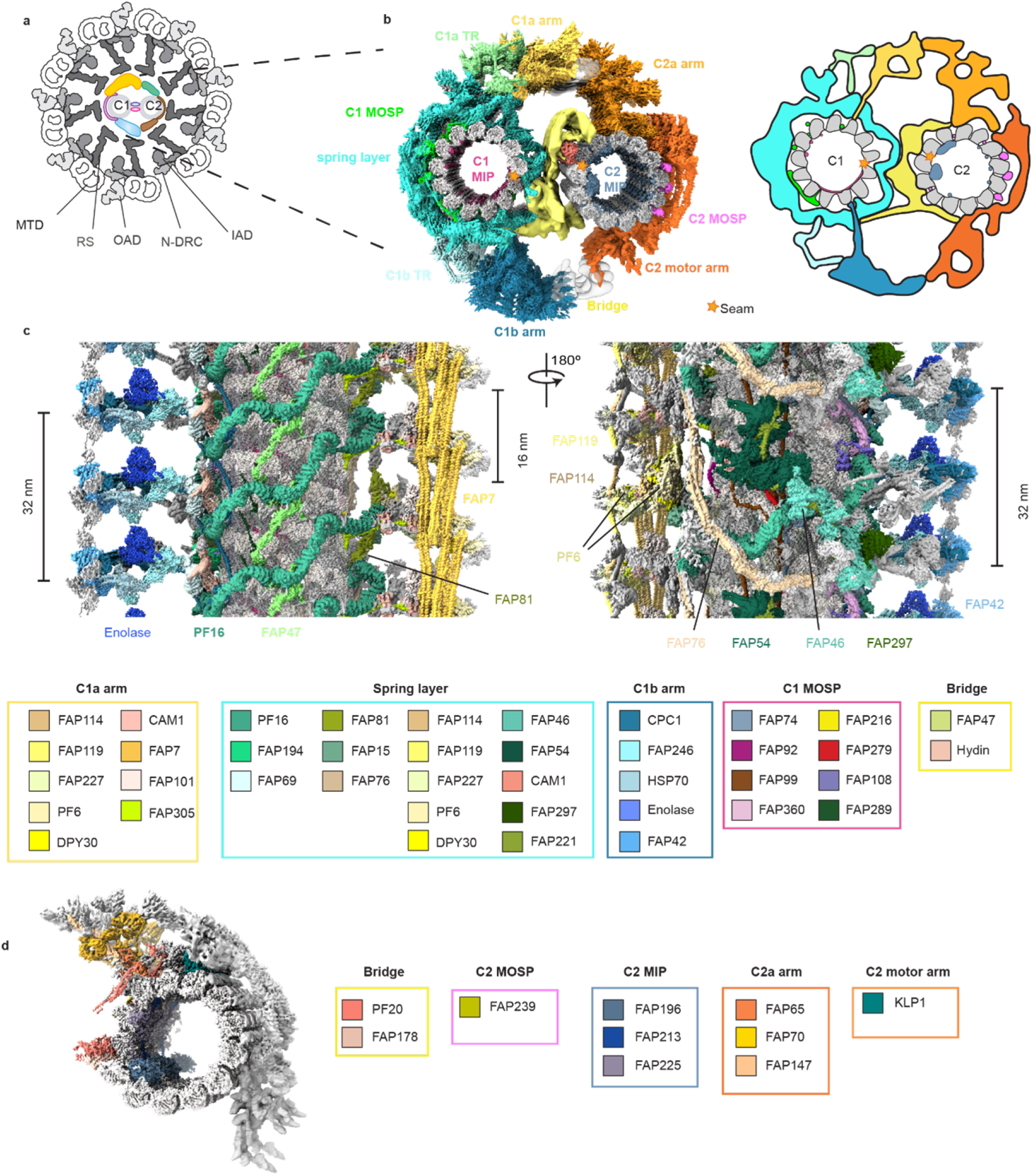
Overview of the central apparatus structure. **(a)** Schematic representation of the cross-section of the *C.reinhardtii* axoneme. The CA in color is surrounded by nine radial spokes (RS) in grey attached to corresponding microtubule doublet (MTD) bound by outer arm dyneins (OAD), inner arm dyneins (IAD), and nexin-dynein regulatory complexes (N-DRC) (13, 15). **(b)** Cross-sectional view of the CA map. CA is divided into 12 regions in different colors: C1 microtubule inner proteins (MIPs), C1 microtubule outer surface proteins (MOSPs), C1 spring layer, C1a transition region, C1a arm, C1b transition region, C1b arm, bridge, C2 MIP, C2 MOSP, C2a arm, and C2 motor arm. A simplified cartoon model is attached on the right. **(c)** Longitudinal views of C1 from two opposite directions. Identified proteins are colored while unassigned density is in grey. Colored components are tabulated below. **(d)** A side view of C2. The components of those regions are listed.

Current knowledge of CA structure is limited to its overall morphology and component characterization. In the past few decades, different models have been proposed to explain how CA functions in ciliary beating (23, 39–41). Nevertheless, due to numerous challenges in obtaining a high-resolution three-dimensional structure of this enormous molecular machine, an in-depth mechanistic understanding of its roles has been largely limited. How the CA components are assembled and work together with each other remains debatable. Furthermore, it is unknown whether CA actively changes its conformations during the beating and why the KLP1 motor is needed for the role of CA.

Here, we determined the cryo-EM structure of a near-complete repeating unit of CA from *C. reinhardtii* at high resolution, providing insight into the assembly of CA and its role in ciliary beating regulation. Our structure shows that functionally related projection proteins of CA are clustered onto a spring-shaped scaffold, which is mainly composed of the armadillo-repeat protein PF16, a homolog of the disease-causing protein sperm-associated antigen 6 (Spag-6) in humans (42–44). One common assembly principle revealed in the high-resolution structure is that each projection complex contains a rachis-like protein that plays a central role to organize all other subunits. The two halves of CA are brought together via a group of flexible, elongated protein complexes that allow relative sliding between the two and may transmit the conformational signals to achieve coordinated activities. Notably, we captured the array of KLP1 in two different stepping states on the C2 microtubule, suggesting that this kinesin plays a role as an active motor system in the central region of a cilium, in addition to the dyneins in the outer regions.

## Results

### Structural determination of CA complex

The complexity and flexibility of CA has limited its resolutions to ~23 Å on C1 alone as the best case reported previously (45).We overcame a series of technical hurdles by optimizing sample preparation, data collection, and image processing to obtain high-resolution cryo-EM structures of CA. To avoid the low contrast by imaging the whole cilium or axoneme, we purified CA by ATP-induced extrusion from the isolated *C. reinhardtii* axoneme. The highly preferred ‘C-shaped’ geometry of CA in cryo-EM grids dictates that ~99% of all segments of CA filaments adopt one single view from the automatically collected cryo-EM datasets. To avoid this extreme orientation problem, we collected high-contrast atlases at low magnification, empirically identified the only ~1% regions that were not severely bent and seemed to display different views by eye, and accurately targeted those regions for the final imaging at high magnification. Due to the variable bending curvatures of CA filament, relative movement between the two halves, and large flexibility of many projections, our attempts for a high-resolution three-dimensional (3D) map using conventional single particle cryo-EM approaches failed, even at the very early stage to generate a usable initial model containing both C1 and C2. A reliable initial model was successfully obtained by cryo-electron tomography (cryo-ET). To tackle the multi-scale flexibility, we utilized a hierarchical local refinement strategy to iteratively focus on individual regions together with re-centering, signal subtraction and local CTF refinement. Using this strategy, we achieved better than 3.5 Å resolutions for the majority of CA by piecing together more than 200 locally refined cryo-EM maps (**Fig. 1b-d, Supplementary Fig. 1, 2**). To avoid possible artifacts around the edges of masked regions during map integration, the masks for each pair of neighboring density maps were created with sufficient overlaps between them. Finally, we built an atomic model of nearly a complete CA by integrating our high-resolution maps with structural information from previous mass spectrometry (46, 47), biochemical characterization (48–52), and mutagenesis studies (49, 53–57). Each 32-nm repeating unit of our final model contains 208 tubulins and 196 non-tubulin chains, which belong to 45 unique proteins and cover ~71% mass of CA (**Fig. 1b-d, Supplementary Fig. 1, 2, Supplementary Table 1, 2**).

### Overview of the central apparatus architecture

CA is an asymmetric complex with distinct projections around C1 and C2 microtubules (29, 30) (**Fig. 1b-d**). All CA sub-complexes are interconnected through intricate networks of scaffolding and connecting proteins. The substantially improved resolutions allow us to re-define the overall organization of CA which was previously divided into C1a-f and C2a-e complexes (29). C1-associated proteins are composed of five layers (microtubule excluded): (1) a mesh of microtubule inner proteins (MIPs), (2) a network of tightly bound microtubule outer surface proteins (MOSPs), (3) a layer of spring-shaped scaffold with associated proteins (spring layer), (4) two transition regions (C1a TR and C1b TR) connecting the spring layer to (5) two large arms (C1a and C1b) projecting towards C2. Similarly, C2 also contains networks of MIPs and MOSPs, as well as two major C2 projection regions, a mobile motor arm sliding on the C2 microtubule and a large stationary arm (**Fig. 1b**). The C1 and C2 microtubules face each other near their seams, brought together by a flexible bridge region (**Fig. 1b**).

### A spring-shaped scaffold wrapping around the C1 microtubule

The flagellar beating is a fast process (~50 Hz in *C. reinhardtii*) (58), during which the CA needs to change its conformations efficiently and drastically without disrupting the structural components. This mechanochemical process necessitates a structural mechanism by which CA can effectively assemble into such a large complex with both structural elasticity and stability. Previous studies revealed that the evolutionarily conserved CA protein PF16 plays critical roles in C1 assembly, stabilization, and ciliary beating (49, 59), of which the human homolog Spag6 has been identified to play emerging roles in many human diseases including cancer (42–44, 60). Our cryo-EM structure reveals that CA contains a layer of spring-shaped scaffold (spring) on the C1 microtubule (**Fig. 2a, b**). The spring contains three highly conserved armadillo-repeat proteins (PF16, FAP194, and FAP69) as basic building blocks. The atomic model shows that the major component of the spring is PF16, which polymerizes on the C1 microtubule surface to form stable spirals (**Fig. 2a,b**).

**Fig. 2:**
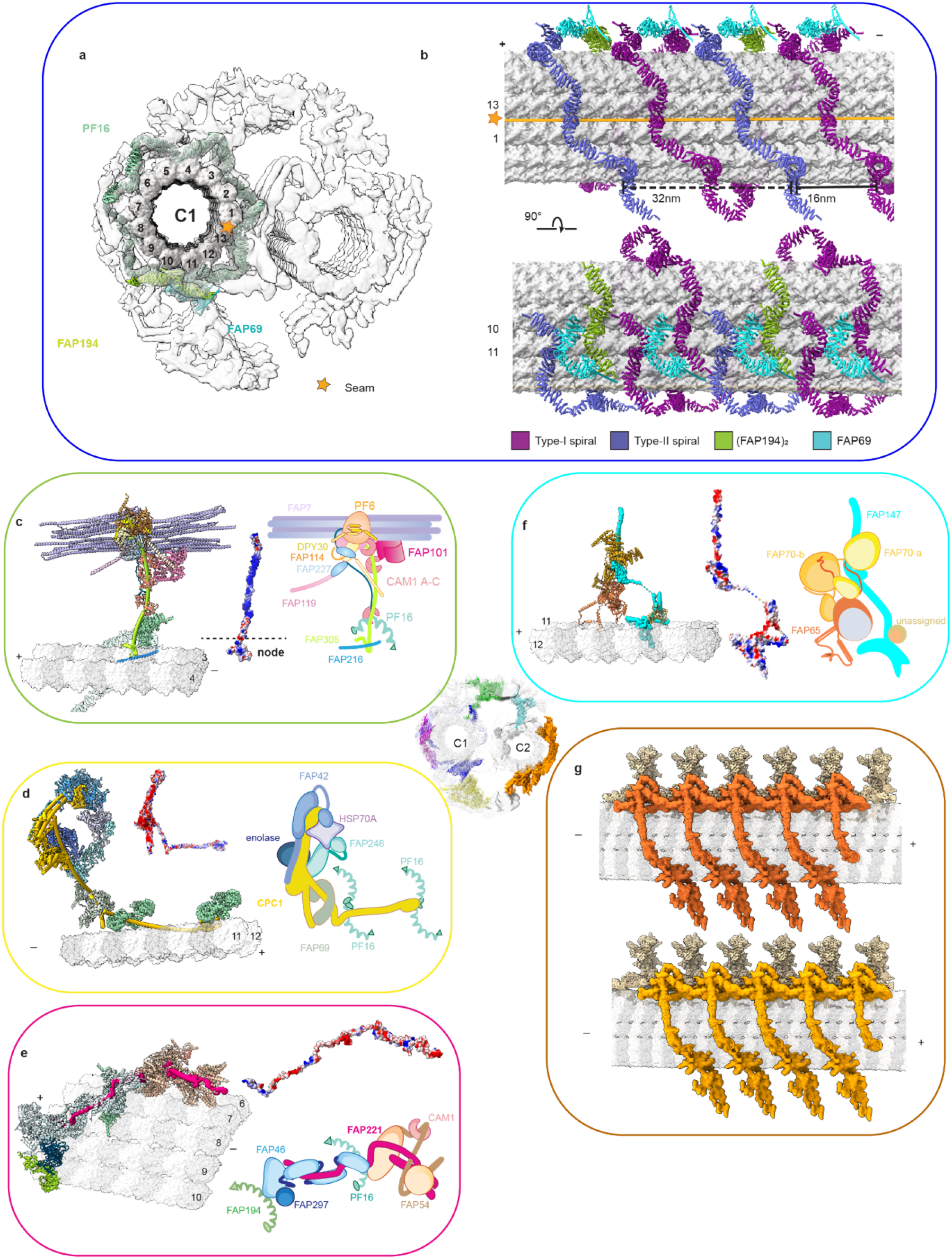
The assembly of CA projections. **(a)** The cross-sectional view of the spring scaffold. **(b)** Longitudinal views of the microtubule-binding pattern of the spring scaffold. Type-I and -II spirals of PF16 are colored purple and blue respectively. One PF194 dimer (green) binds to the protofilament 10. In each 32-nm repeat, one FAP69 monomer (cyan) seals the termini of neighboring type-I spirals while the other FAP69 monomer connects the FAP194 dimer to type-II spirals. **(c)** The structure of C1a arm. The atomic model (left) of C1a arm contains the rachis protein FAP305 (green cylinder), Calmodulin 1 (CAM1), FAP7 and the PF6 complex (composed of PF6, FAP114, FAP119, 2x FAP227, and 2x DPY30). The electrostatic map (middle) of FAP305 shows charged patches. A cartoon model (right) of C1a arm is attached. **(d)** The structure of C1b arm. The atomic model (left) of C1b arm contains rachis protein CPC1 (yellow cylinder), FAP246, FAP42, HSP70A, and Enolase. The electrostatic map (middle) of CPC1. A cartoon model (the) of C1b arm. **(e)** The structure of C1d complex. The atomic model (left) of C1d complex contains the rachis protein FAP221 (pink cylinder), CAM1, FAP54, FAP46 and FAP297. The electrostatic map (middle) of FAP221. A cartoon model of C1d complex is attached on the right. **(f)** The structure C2a arm. The atomic model (left) of C2a arm contains rachis protein FAP147 (in cyan cylinder), FAP65, FAP70, and an unassigned protein. The electrostatic map (middle) of FAP147. A cartoon model (right) of C2a arm. **(g)** Overviews of the two classes of C2 projections. The motor arm (orange and dark orange) slides on the C2 microtubule, while C2a arm (light yellow) is immobile. The map in the center shows the locations of the six representative projections of CA, including spring scaffold (blue), C1a arm (green), C1d complex (pink), C1b arm (yellow), C2a arm (cyan), and motor arm (orange).

PF16 contains 11 consecutive armadillo motifs and additional helices at both N- and C-termini (**Supplementary Fig. 3a**). The armadillo repeats form a right-handed alpha-solenoid (**Supplementary Fig. 3b**), with strikingly polarized charge distribution (**Supplementary Fig. 3c**). Each positively charged alpha-solenoid groove of PF16 clenches a tentacle-like C-terminal tail of α-tubulin (**Supplementary Fig. 3d-g**) and harbors an elongated region of a CA projection protein (**Supplementary Fig. 3e, h**) which is linked to the rest of CA projections. By contrast, the outer surface of PF16 carries negatively charged residues for binding positively charged CA proteins.

PF16 molecules dimerize at their N-termini (**Supplementary Fig. 3i, j**). PF16 dimers further polymerize on the C1 microtubule surface via their C-termini to form vine-like spirals. Each 32-nm repeat of CA contains two types of spirals termed type-I and -II (**Fig. 2b**). The type-I spiral contacts every other microtubule protofilaments and wraps around C1 microtubule surface for a complete turn. The type-II spiral interdigitates with type-I spiral at a 16-nm interval, but only forms a half turn. Proteins made of armadillo repeats usually display high flexibility in tertiary structure, whereas their secondary structure elements remain stable in response to external tension, similar to that of a mechanical spring (61). We compared the structures of all the 20 subunits of PF16 from one repeating unit of CA and observed large structural variations among them, while their secondary structures are preserved to very high fidelity (**Supplementary Fig. 4a, b, Supplementary Video 1**). To understand why PF16 displays a variety of elastic conformational changes on the CA microtubule, we analyzed the rotation angles between each pair of adjacent microtubule protofilaments (inter-pf angles). Notably, the inter-pf angles of both microtubules of CA vary substantially (**Supplementary Fig. 4c, d**), distinct from that of the cytoplasmic microtubule which is approximately 13-fold symmetric. The variable inter-pf angles of CA microtubules require different distances among the PF16 binding sites at different locations on the microtubule surfaces. Strikingly, the PF16 molecules can elastically change their tertiary structures to adapt to such geometric changes without breaking their secondary structures, suggesting the spring scaffold has a potential to behave like a mechanical spring to furnish CA with the structural elasticity and stability, which is essential for ciliary beating.

In addition to PF16, the spring scaffold also contains two other armadillo-repeat proteins, FAP194 and FAP69 (**Fig. 2a, b, Supplementary Fig. 3k**). FAP194 is highly similar to PF16 in its sequence, structure, dimerization interfaces, and interactions with microtubule, but has a longer N-terminal helix. FAP69 functions as monomers to join the termini of two adjacent type-I spirals or connect type-II spirals to FAP194. In addition, the spring scaffold is tied to the network of peptide-like MOSPs, which in turn penetrate through the clefts between adjacent microtubule protofilaments to interact with the MIPs (**Supplementary Fig. 5**). Thus, all elements of the spring scaffold as well as microtubule outer and inner surface proteins are joined together to form an interconnected network, which stabilizes and elasticizes the C1 microtubule. Interaction analysis reveals that the spring scaffold also serves as a hub for organizing all other CA projection proteins (**Supplementary Fig. 6**), explaining the essential role of PF16 in CA assembly.

### General assembly principle of CA projections

CA projections extend out from the microtubules and the spring layer, branching into smaller sub-complexes, like a cluster of flowers. This principle governs the organization of at least four major projections, including C1a arm, C1b arm, C1d complex, and C2a arm. Central to each of the four projections is a unique rachis-like protein that clusters functionally related projection proteins. Each large projection is organized as smaller sub-complexes branching out of the rachis (**Supplementary Fig. 6**). All CA projections are interconnected via elongated connecting proteins that define the degrees of relative movement among different regions. Nearly every CA projection is associated with one or more EF-hand proteins, which have the potential to change their conformations upon calcium-binding (52, 62–64), raising the possibility that the behavior of CA can be regulated by the calcium influx through the conformational changes of those EF-hand motifs.

### Assembly of C1 projections and their potential interactions with radial spokes

C1a arm plays an important role in the transmission of mechanical feedback from CA to radial spokes via electrostatic repulsion (65). Assembly of C1a arm centers around the rachis protein FAP305, which stems out from the C1 microtubule protofilament-3, attaches to the spring scaffold and clusters all other C1a subunits and sub-complexes (**Fig. 2c**). C1a arm contains a compact sub-complex which we name PF6 complex (containing PF6, FAP114, FAP119, 2x FAP227, and 2x DPY30). Another copy of PF6 complex is also found in the C1e region (**Supplementary Fig. 7**). PF6 is a highly conserved protein required for C1a assembly and was proposed to mediate radial spokes-CA interactions (35, 49, 54). The two PF6 complexes are both located at the outermost surface of CA, separated by approximately the same spacing between two adjacent radial spokes (**Supplementary Fig. 8a**), which allows a pairwise radial spoke-PF6 geometry match for direct contacts. The surfaces of PF6 complexes and radial spokes at the interfaces are all negatively charged (**Supplementary Fig. 8b, c**), which strongly supports a previously proposed model (65) that electrostatic repulsion occurs when they are close enough during the beating.

C1b arm was proposed to regulate the ciliary beating frequency through nucleotide concentration maintenance (50, 57). Knock-out of the C1b component CPC1 leads to the loss of entire C1b arm in *C. reinhardtii* (53), and mutations of the mammalian homolog of CPC1 lead to immotile sperms and primary ciliary dyskinesia (PCD) (66, 67). Our structure shows that CPC1 serves as the rachis of the entire C1b arm. CPC1 is a large, elongated protein that spans along the C1 microtubule longitudinally and also extends out perpendicularly for over 20 nm. The extended CPC1 organizes a large cluster of C1b projection proteins with various enzymatic activities for energy metabolism, including FAP246, FAP42, HSP70, and enolase (**Fig. 2d**). In addition, CPC1 stabilizes C1b arm by laying on C1 microtubule protofilaments to connect adjacent PF16 spirals of the spring scaffold. Besides its role as the C1b rachis, CPC1 also has an adenylate kinase domain for potential ATP level regulation and an EF-hand domain for sensing calcium signals.

All the C1b proteins clustered by CPC1 contain enzymatic domains for controlling nucleotide levels collectively (50), such as the guanylate kinase domain and EH-hand pair in FAP246, and the cysteine peptidase domain, adenylate kinase domain, and four guanylate kinase domains in FAP42. Another protein we identified in C1b arm is enolase, which catalyzes an essential reaction in the glycolysis pathway for ATP production (50). All these enzymatic domains are spatially close to each other, highlighting the role of C1b arm as a potential ‘chemical factory’ for coordinated activities to control the distribution of nucleotides required for beating regulation. On the other hand, the surface of C1b arm that faces the radial spokes is also negatively charged, which provides another repulsion site between CA and radial spokes (**Supplementary Fig. 8d**). C1d projection affects the waveform and speed of *C. reinhardtii* (33, 62, 68) and is implicated in PCD (69). C1d contains a rachis protein FAP221, which clusters FAP46, FAP54, FAP297 and CAM1 (**Fig. 2e**). C1d is predominantly composed of helical domains which together form an extended layer and attach to the top of the spring scaffold at multiple sites (**Supplementary Fig. 8a**), potentially reinforcing the elasticity and stability of the spring. C1d surface is not obviously charged (**Supplementary Fig. 8e**) but may still provide additional interaction sites with two adjacent radial spokes for the perfect geometry match.

### C2 contains stationary and motile projections

C2 projections display features distinctive from those of C1, which lead to asymmetric CA projections for differentiated roles of the two halves. In contrast to C1 projections with fixed axial positions along the C1 microtubule, C2 projections contain both stationary (C2a arm) and motile regions (motor arm). C2a arm is assembled around a rachis protein FAP147, which organizes other C2a proteins (e.g., FAP65 and FAP70) (**Fig. 2f**). Notably, we identified a motile region that includes the C2b, C2c, C2d and C2e described in previous study (29). We found this region can slide on the C2 microtubule up to 8 nm. Extensive 3D classification suggested all sub-complexes in this region move together along the microtubule, despite the co-existence of severe local structural flexibility. Therefore, we name this region the CA motor arm to reflect this conformational behavior. Particularly, two different locations and states of the motor arm are observed by focused cryo-EM classification (**Fig. 2g**). In both states, the motor arm is held onto the C2 microtubule through the motor protein KLP1 at one end (adjacent to C2a arm) and linked to C1b arm at the other end. Proteins on the motor arm are extremely flexible and only loosely contact the C2 microtubule surface, distinctive from those tightly bound CA projections. Such features of the motor arm enable its sliding on the microtubule track more freely with less energy consumption (Supplementary Video 2). The dynamic motor arm is located right next to the ‘chemical factory’, raising the possibility that the movement of the motor arm may be primarily supported by the newly synthesized ATP molecules from C1b arm.

### KLP1 forms an active motor array on the C2 microtubule

The kinesin KLP1 was suggested as a component of C2 projections (48). Its knockdown hampered the normal flagellar motility in *C. reinhardtii* (23), and mutations of its mammalian homolog Kif9 severely affected sperm motility (70). However, it is unknown whether KLP1 is active and how its mutations impact ciliary motility. Our cryo-EM and cryo-ET structures of the KLP1 array on C2 in different states provide key evidence for its role as an active motor in cilia.

KLP1 forms an asymmetric dimer with both the leading and trailing heads bound to C2 protofilament-9. The dimers further assemble into an array mediated by their tail domains and associated proteins (**Fig. 3a**). The trailing head of KLP1 adopts the so-called neck-linker docking conformation (**Supplementary Fig. 9**), whereas the leading head is in an undocked conformation (**Fig. 3b**). The neck-linker conformation typically represents an ATP-bound state in free kinesins in vitro (71), but our structure of the native KLP1 in the array contains an ADP instead (**Fig. 3b, Supplementary Fig. 9b**). As the CA sample was pre-treated with ATP during purification, the conformation captured in this study is most likely an intermediate state after ATP hydrolysis and before ADP release. Intriguingly, we used the isolated CA after ATP treatment to find that KLP1 array adopts two different microtubule-binding states (MTBS-1 and −2) on the C2 microtubule. From MTBS-1 to MTBS-2, the trailing heads of the KLP1 array cooperatively move 16 nm forward, while the leading heads preserve their original positions on the C2 microtubule. The movement observed in our cryo-EM structure agrees with previously proposed ‘hand-over-hand’ model (72) on free-form kinesins. The conformational changes of the head regions of KLP1 array result in an 8-nm movement of the KLP1 tail complexes, which finally drives an 8-nm sliding of the entire motor arm on the C2 microtubule (**Fig. 3c**).

**Fig. 3:**
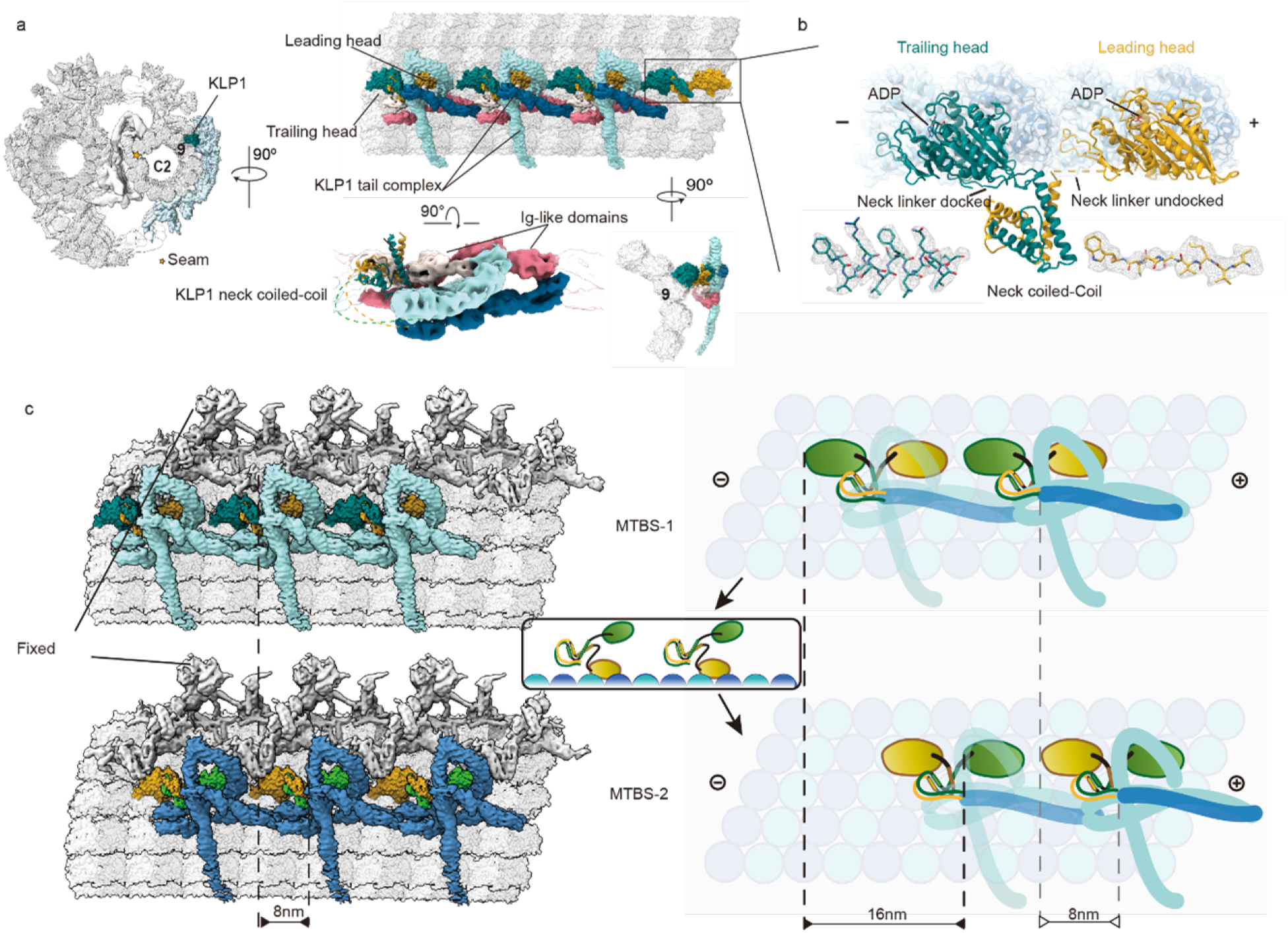
Two microtubule-binding states of the KLP1 array. **(a)** The kinesin-like protein 1 (KLP1) lines up in an array on the C2 microtubule protofilament 9. The leading head and trailing head are colored gold and dark cyan respectively. KLP1 tail complex contains several consecutive Ig-like domains. **(b)** The structure of dimeric KLP1. KLP1 dimerizes around the necks, while the two heads, both containing ADP, separately bind to the microtubule. The trailing head adopts the neck-linker docking conformation, whereas the leading head is in an undocked conformation. **(c)** The mechanism of the motor arm movement. For MTBS1 to MTBS2, the KLP1-associated proteins move 8 nm forward driven by the trailing head that steps 16 nm to the plus end, while the leading head stays stationary with respect to C2a arm. The rest regions of the motor arm move along with KLP1-associated protein as they together form an interconnected complex.

### Elastic C1-C2 connections for conformational communication

C1 and C2 are held together at three major regions, a central bridge connecting the two microtubules and two flanking links (link-a and link-b) between the opposing C1 and C2 arms at the periphery (**Fig. 4a**). We identified four proteins (hydin, PF20, FAP47, and FAP178) in the bridge, together forming chain-like connections between C1 and C2. The chains are mainly constituted by consecutive Ig-like folds from hydin and FAP47 (**Fig. 4b**). The hydrocephaluscausing CA protein hydin was previously proposed to localize on C2 (73). Interestingly, our high-resolution structure shows that hydin also anchors onto the spring scaffold of C1 with its N-terminus tightly bound to the C1 microtubule, albeit the majority resides on the motor arm (**Fig. 4c**). Similarly, FAP47 was thought to be a C2 component (47) but we find its N-terminal region also tightly binds to C1 (near the seam) and links up adjacent spirals of the spring scaffold on C1 (**Fig. 4d**). The C-terminal region of FAP47 is too flexible to allow atomic model building. Nevertheless, the weakly connected density suggests that FAP47 extends towards the C2 microtubule and potentially binds other bridge proteins to form a complex (47) which we name the FAP47 complex (**Fig. 4b**). The complex interacts with PF20, an essential protein for the assembly of entire CA and ciliary motility (51). Our structure suggests PF20 plays a central role by holding the two halves of CA together. The N-terminal helix of PF20 dimerizes under the base of C2a arm and stabilizes the binding between C2a arm and the C2 microtubule. The C-terminal WD40 domain of PF20 interacts with FAP178 to form a complex which lays across the C2 protofilament-1 and −13 to button up the seam (**Fig. 4e**). The PF20/FAP178 complex also contacts the FAP47 complex which in turn is connected to the C1 microtubule seam. We therefore speculate that PF20 plays a critical role, together with the FAP47 complex, in arranging the relative positions of the C1 and C2 seams, which may explain why PF20 is essential for CA assembly (51).

**Fig. 4:**
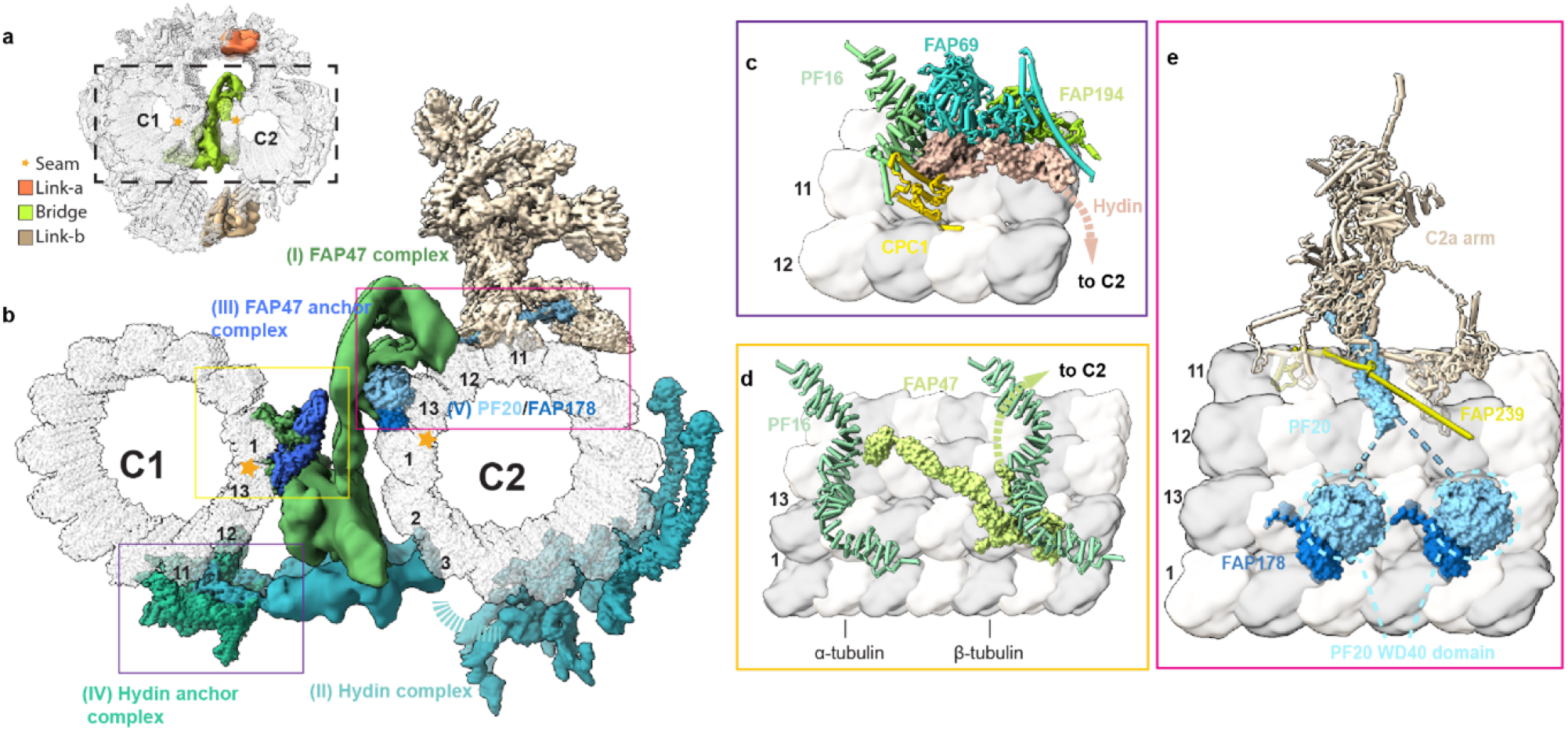
The connections of two halves of CA. **(a)** An overview of three contact sites between C1 and C2. Link-a (orange) connects C1a arm and C2a arm, while link-b (wheat) connects C1b arm and C2 motor arm. The bridge (green) directly connects two microtubules. **(b)** An overview of the bridge architecture. The bridge is divided into five parts: the FAP47 complex (I) and Hydin complex (II), which together form the main structures mediating the connection of two microtubules, the FAP47 anchor complex (III), Hydin anchor complex (IV), and the PF20/FAP178 complex (V). **(c-e)** Atomic models of three microtubule-bridge association sites. Hydin is anchored onto the C1 microtubule protofilament 11 at the base of C1b arm, while the C-terminal region is located on the C2 motor arm **(c)**. The N-terminus of FAP47 links two PF16 dimers from adjacent spirals and binds to the C1 microtubule protofilament 13 and 1 while the C-terminal region extends toward the C2 microtubule **(d)**. PF20 dimerizes with its N-terminal long helix under the base of C2a arm while its C-terminal WD40 domain (dashed circle) forms a complex with FAP178 on the C2 microtubule seam **(e)**.

A notable feature of the chain-like structures in the bridge is their flexibility and elasticity, which eventually defines the extents of relative sliding and rotations between the two halves for conformational communications. By aligning the repeating units from the same CA, we find the two halves continuously slide against each other in the curved regions of CA. The range of relative sliding is continuously distributed from −12 nm to +12 nm longitudinally with two notable peaks distanced by 8 nm (**Supplementary Fig. 10a**), which is precisely the step distance of KLP1 on the C2 microtubule, implying that such distribution is correlated with the microtubule-binding states of this kinesin motors. In addition to the sliding, C1 and C2 also twist against each other. The amount of relative twist is usually small (within 2 degrees) but there is a small ratio (~ 10%) of CA segments that can undergo significant twists (up to ~10 degrees) (**Supplementary Fig. 10b**). To further understand how these geometry parameters affect the overall conformations of CA, we performed cryo-ET reconstructions of CA with different shapes. The analysis reveals that C1 is always on the convex surface of CA and bends towards C2 (**Supplementary Fig. 11a**), suggesting that there exist accumulative tensions between the two halves along CA. We estimated the relative C1-C2 twists between adjacent repeating units. In line with our single particle cryo-EM analysis, the twists in most regions are relatively small except the relatively straight regions where the bending phases of CA are inverted along with a sharp twist between C1 and C2 (**Supplementary Fig. 11b**). The maximum amount of twist also agrees with the estimated distribution of relative rotations between C1 and C2 by single particle analysis (**Supplementary Fig. 10b**). The sharp twist in the phase-transition region necessitates large structural changes around the bridge region. These geometry changes are ultimately defined by the movement of KLP1 arrays, whose mechanical forces are transmitted from C2 to C1 through the chain-like bridge proteins and eventually affect interactions between CA and radial spokes.

## Discussion

The beating of eukaryotic cilia and flagella is a rhythmic process with radially asymmetric motor activities (33, 35, 36, 74), which are critical for directional movement and energy efficiency. Nearly all motile cilia require the CA as an indispensable core component and the loss of CA leads to paralyzed cilia (25, 30, 35, 53). Our high-resolution cryo-EM structure leads to a substantially improved model of CA architecture (**Supplementary Fig. 12**). The microtubule inner and outer surface proteins serve as binding wires to stabilize the core of CA and define the repeating units of CA. The spring scaffold, mainly composed of the armadillo-repeat protein PF16, wraps around the C1 microtubule to function as an assembly hub for all other projections. The ability of PF16 to elastically change its conformation with high fidelity of secondary structure suggests that the spring scaffold has a potential to endow CA with structural elasticity required for beating. Each large CA projection has an elongated rachis-like protein that anchors onto the spring scaffold and clusters functionally related proteins. The C1a arm and the C1e region, each containing the negatively charged PF6 complex at the convex surface of CA, provide a collision center for electrostatic repulsion between CA and radial spokes. C1b arm is composed of several enzymatic domains and forms a ‘chemical factory’ for maintaining the nucleotide level in cilia. C2 projections are generally more dynamic due to the flexibility of the motor arm. The KLP1 array on C2 powers the movement of the motor arm, which in turn leads to active geometry changes of the whole CA for ciliary beating regulation. The chain-like bridge proteins transmit the mechanical forces generated by the motor arm to C1 for coordinated conformational changes between the two halves of CA.

Furthermore, we docked our high-resolution structure of *C. reinhardtii* CA to several previously reported cryo-ET maps from different species (29, 75) and found both the spring layer and major projections fit very well (**Supplementary Fig. 13a-d**), suggesting structural and functional conservations across species. To locate ciliopathies-related proteins, we built homology models of human CA projections (**Supplementary Material 1**) and compared with those of *C. reinhardtii* CA determined in our study. A notable finding is that nearly all the CA proteins identified from our cryo-EM maps (37 out of 45, excluding tubulins) have close human homologs with most of the CA projection cores more conserved (**Supplementary Fig. 13e**). Therefore, the CA structures we determined will also provide rich information to guide future studies on the roles of human CA in ciliopathies.

## Materials and methods

### Strains and cell culture

*Chlamydomonas reinhardtii* wild-type strain (CC-124) was obtained from the Chlamydomonas Resource Center (https://www.chlamycollection.org). Cells were grown in TAP (Tris-acetate-phosphate) medium under continuous aeration and illumination for three days at 25 °C to reach a cell density of ~2.5-4 × 10^7^ cell/ml (OD750 ~3).

### CA isolation

We used a previously published protocol (76, 77) with some modifications to isolate CA. The flagella were excised from *C.reinhardtii* cell bodies by dibucaine treatment at a final concentration of 2.5 mM for 2 min. The purified flagella were resuspended into HMDEKP buffer (30 mM HEPES, 5 mM MgSO4, 1 mM DTT, 0.5 mM EGTA, 25 mM KCl, 1 mM PMSF, pH 7.4) with 1% IGEPAL CA-630 on ice for 10 min to solubilize flagellar membrane. The ‘9+2’ axonemes were centrifuged at 20,000 g for 10 min. The pellets were resuspended in HMDEKP buffer and incubated with 3 mM ATP at 25 °C for 1 hour. After the reaction, 10 mM ATP was added to the axoneme. The axoneme was incubated for 1 hour at 15 °C and then centrifuged at 6,000 g for 5 min to remove the ‘9+0’ axoneme. The supernatant, which mainly contained the CA and split microtubule doublet, was pelleted by centrifugation at 20,000 g for 10 min and then resuspended at a ratio of 20 μL HMDEPK buffer per liter cultured cells for cryo-EM analysis.

### Cryo-EM sample preparation

Cryo-EM grids of CA were prepared using Vitrobot Mark IV (Thermo Fisher Scientific). Three microliters of the purified CA sample were applied to each Quantifoil holy carbon grid (R2/1, 300 mesh gold). All grids were incubated for 5 s in the Vitrobot chamber at 8 °C and 100% humidity, blotted with standard Vitrobot filter paper (Ted Pella, Inc.), and then plunged into liquid ethane at approximately −170 °C.

### Cryo-EM data collection

All data collection was automated by SerialEM software (78). The Dataset 1 (**Supplementary Table 1**) was collected on a 300 kV Titan Krios microscope (Thermo Fisher Scientific) equipped with a Bioquantum Energy Filter and a K2 Summit direct electron detector (Gatan) at Yale CCMI Electron Microscopy Facility. 10,047 movies were recorded using beam-tilt induced image-shift protocol (5 images for each stage movement) at a total dose of 39.2 e-/Å^2^ per movie and a defocus range from −1.2 to −2.5 μm. The nominal magnification was originally set at 130,000 x, corresponding to a calibrated pixel size of 1.05 Å at the super-resolution mode (0.525 Å per super-resolution pixel). After preliminary 2D analysis, we found that nearly 99% of the filament segments of CA from the fully automatically collected data fell into a single class, due to the severe propensity of CA to adopt a ‘C-shaped’ geometry in our cryo-EM grids. The ‘C-shape’ corresponds to a conformational state in which the C1 half is always on the convex side of CA, bending towards the C2. By careful analysis of the relatively straight regions, we found those ~1% particles provided more orientations for a promising 3D reconstruction. We therefore increased the pixel size to 1.33 Å for the following two datasets as a compromise between the achievable resolution and effective particle number. To further overcome the preferred orientation of CA, we collected a set of atlases for all promising squares in the view-mode at ~4 times longer exposure than that of normal single-particle data collection, identified each CA by eye, guessed its orientation empirically, and then manually assigned each individual target for the final high-magnification data collection. The Dataset 2 (**Supplementary Table 1**) containing 3,060 movies was collected at Yale CCMI Electron Microscopy Facility using a similar setting except that the nominal magnification was 105,000 x and the total dose per movie was increased to 48.4 e-/Å^2^. At this stage, the number of particles still remained a major limiting factor for high-resolution structure determination. Therefore, a third dataset (Dataset 3 in **Supplementary Table 1**) was collected on a 300 kV Titan Krios microscope (Thermo Fisher Scientific) equipped with a K3 detector (Gatan) and a Bioquantum Energy Filter at Case Western Reserve University Electron Microscopy Facility. We used a nominal magnification of 65,000 x, corresponding to a calibrated pixel size of 1.33 Å, at a total dose of 38.6 e-/Å^2^ and a defocus range of −1.2 to −2.5 μm. 5,175 movies were acquired at the correlated double sampling (CDS) mode. All three datasets were scaled to 1.33 Å for reconstruction.

### Cryo-EM Data processing

For all datasets, motion correction was performed by MotionCor2 (79), CTF was estimated by Gctf (80), and particles were picked by Gautomatch (https://github.com/JackZhang-Lab). The processing was streamlined using a modified script available at https://github.com/JackZhang-Lab/EM-scripts. Particles were extracted from the dose-weighted micrographs in RELION V3.0 (81) and imported into cryoSPARC v2 (82) for all subsequent processing.

At the very beginning, particles were automatically selected by Gautomatch using a Guassian blob as the template. The top 20 classes averages were used as the new templates for subsequent auto-picking. The templates were updated once micrographs had been collected to improve the accuracy. A total number of 1,381,963 particles were automatically selected by Gautomatch with a 12-nm distance cutoff (the one with a lower cross-correlation was rejected if the distance from another one is less than 12 nm). To speed up the data processing, all particles were downscaled to a pixel size of 5.32 Å and a box size of 256*256. After several cycles of 2D classification, 359,888 high-quality particles were selected for the final refinement and 3D classification. Even though we had limited the data collection to relatively straight regions of the CA filaments, extensive 2D classification suggested that a wide range of bending curvatures still existed in the final datasets collected. The various bending curvatures are correlated with relative slides and twists between the two halves of CA (**Supplementary Fig. 10–11 and Supplementary Video 2**). In addition, many of the projections themselves have large local flexibility as well as relative movements among them. For these reasons, we failed to obtain a usable initial model containing both C1 and C2 by the singleparticle approach in cryoSPARC (82) or Relion (81). A reliable initial model was successfully obtained by refining a previously reported cryo-ET map EMD-5853 (35) using subtomogram averaging approach in IMOD (83, 84).

Due to the multi-scale flexibility of the CA structure and severely overlapped signals from such an enormous target, local refinement by simply providing a set of masks in different regions failed to improve the resolutions. This issue could not be solved by providing a wider range of parameter search during local refinement, which would immediately lead to a divergent parameter estimation. For instance, the sliding between C1 and C2 is up to about 120Å, which leads to a complete loss of interpretable C2 density if we used the C1 parameters to reconstruct C2. To deal with this issue, we utilized a hierarchical local refinement strategy to gradually focus on individual regions together with re-centering, signal subtraction, and local CTF refinement (**Supplementary Fig. 1**). We first divided the whole complex into multiple regions at several different levels. The first level corresponded to the two halves of C1 and C2. The second level represents the microtubules together with surface-binding proteins and projection complexes. The third level covers sub-complexes of the projections and 3-5 protofilaments with their binding proteins within each mask. Beyond the third level, we further divided those local regions into smaller ones for additional cycles of local refinement.

For the local refinement at the first three levels, our overall strategy is to iteratively improve the estimated centers for each target region against individual particles and iteratively remove the background by signal subtraction. Once we had an initial reconstruction of C1, we removed the C1 signal from the particles, pasted the subtracted particles back to their original positions, and re-extracted the C2 particles free of C1 signals. This in turn generated a better C2 map, which was subtracted from the original particles to improve C1. The approach was performed over cycles to iteratively improve the density maps of both halves along with the improvement in the alignment parameters. During the iterative refinement, we also utilized the priors of the CA conformational changes to guide the settings of the local refinement parameters. For instance, we learned from the 2D analysis that there existed a sliding between C1 and C2, and therefore restricted the search of alignment parameters mainly along the C2 microtubule axis, instead of a generally wider range of parameter search. The initial parameters of individual projections were estimated in a similar way. Eventually, we generated individual datasets that were re-centered and reextracted to focus on each local region with a relatively clean background by signal subtraction.

The metadata format conversion was performed by the UCSF script Pyem (https://github.com/asarnow/pyem) to facilitate particle subtraction, re-centering, and reextraction. Particles were re-centered and re-extracted to a pixel size of 2.66 Å and a box size of 512*512 pixels for the overall structure of C1. To determine the C1 structure within a 32-nm repeat, 3D classification of the selected particle was performed to produce two different classes that displayed a 16-nm shift between them. After classification, all selected particles were re-centered and re-extracted with a pixel size of 1.33 Å and a box size of 832*832 pixels for subsequent processing.

The class containing 190,727 particles was used for subsequent processing to refine the C1 structure. The entire C1 volume was generated by a local refinement with a mask on C1. After that, we re-centered the particles, reduced the box size, and performed signal subtraction for different regions to improve their resolutions. For the microtubule and its surface binding proteins, we carefully re-centered the particles at C1 microtubule and reduced the box size to 512*512 pixels to achieve higher resolution. Subsequently, a series of smaller masks that cover *sim*2×4 tubulin lattice and associated structures were applied to improve the quality of the density map. To further improve the alignment accuracy of a certain projection, the signals from CA microtubules and other binding proteins were subtracted from raw particles.

For the C2 structure determination, C1 signals were subtracted from all selected particles. Several rounds of 2D classification and 3D classification were performed to select high-quality particles from the subtracted data. Totally, 192,253 particles were selected for subsequent local refinement on C2 using the same strategy as described for C1. As the motor arm is too flexible, 3D classification was performed on the particles with the microtubule signals subtracted.

All locally refined maps were aligned and stitched together to generate density maps of the two halves of CA in Chimera using the command vop maximum (85).

### Identification of CA proteins

45 unique non-tubulin proteins of CA were identified by combining previously published mass spectrometry data (46, 47), structure prediction by Phyre2, and sequence pattern search using own scripts (86). First, each of the subunits that show clear backbones was manually built as a poly-Ala model in the local regions with the best resolutions in Coot (87, 88). All side chains were tentatively divided into several groups: (1) large (Trp, Try, Arg, Phe, His); (2) middle (Leu, Gln, Asn, Ile, Met, Lys); (3) small (Pro, Val, Ser, Thr, Cys, Glu, Asp, Ala); and (5) Gly. To improve the identification accuracy, we only used the regions that we were fully confident with the backbone assignment and avoided using the ambiguous regions. A sequence pattern was tentatively generated according to side chain density and was used to search against the CA sequence database. The database was based on CA proteomic mass spectrometry results and the Chlamydomonas Flagellar Proteome Project (89). The search was based on a Linux ‘gawk’ script (https://github.com/JackZhang-Lab) that aims to match a user-defined regular expression pattern, which was regarded as a ‘fingerprint’ of the target. In practice, the pattern search was performed semi-automatically, and the user-defined pattern was iteratively improved upon the results of the hit candidates. For example, if the output contains too many protein sequences, we would either use a stricter role for the group assignment of the residues with high-quality side chains or use a longer peptide to decrease the number of false hits. By contrast, if there is no output using a certain pattern, there must be errors in the search pattern, or the candidate database does not include the target sequence. In this case, we would relax the group assignment to allow more flexibilities or use a larger database, such as the ciliary database or the entire proteome of *C. reinhardtii* (*Creinhardtii*_281_v5.6 protein). Multiple regions were attempted for identified proteins to mutually verify the results with each other. We discarded those hits that conflicted with the hits using different regions from the same protein. The majority of CA projection candidates that match the sequence pattern from our cryo-EM map could be found in the CA database except FAP213 and FAP196, which were further searched out from the whole Chlamydomonas proteome database. We manually checked all side chains of every candidate to verify whether the sequence matches the density. We also excluded the possibility of other homologs for each identified protein by simple side-chain substitution. A protein is defined as ‘identified’ only if the assigned sequence best matches the cryo-EM density without other possibilities in the whole proteome database.

### Model Building and Refinement

Different approaches were used to build models depending on the resolutions. Most of the regions were refined at better than 3.5 Å resolution, which allowed us to build atomic models with side chains assigned in Coot (87, 88). For the slightly worse regions, we were able to build backbone models with the residues assigned based on the relative positions among the large residues (such as Try and Arg) of each domain. For the regions that showed a clear backbone with low-quality side chain density, we coarsely assigned the residues using the predicted domain model from Phyre2. For those regions that were solved at a resolution with clear secondary structures, we fitted the predicted models into the density as rigid bodies in Chimera (85). All models at better than 4 Å resolution were automatically refined by Refmac5 (90) or Phenix (91), and manually checked in Coot. The process was repeated until all parameters were reasonably refined. The figures and movies were created by Chimera (85), ChimeraX (92), and FIJI (93).

### Cryo-ET data collection and tomogram reconstruction

Tomographic datasets which contained an S-shaped CA were collected on the 300 kV Titan Krios equipped with a K2 detector at Yale CCMI Electron Microscopy Facility. SerialEM (78) was used for automatic data collection under the bidirectional scheme at a 2 ° interval and tilt angles ranging from −60 ° and +60 °. Each of the final tilt series contains 61 movie stacks with a pixel size of 5.4 Å, defocus at −5 μm, and an accumulative dose of 120 e-/Å^2^. The recorded movies were motion-corrected using MotionCorr2 (79). The tomograms were reconstructed by the IMOD software package (83, 84).

### Sub-tomogram averaging and reverse fitting

After tomogram generation, C1 and C2 particles were manually picked in a start-to-end manner. The function of addModPts from PEET (15) was used to generate 8-nm microtubule particles. Subsequent sub-tomogram averaging was performed on C1 and C2 separately. Using a 13-protofilament microtubule as a reference map together with a cylinder mask, these particles were aligned and shifted to the center of the microtubule after alignment. Then the original coordinates were re-centered using the parameters from the sub-tomogram analysis. We performed polynomial fitting with degree three for the re-centered coordinates and generated the new particles with 32-nm spacing and corresponding Euler angles (2 out of 3). Those particles with 32-nm spacing in C1 and C2 were iteratively aligned to the low-pass (80 Å) filtered cryo-EM maps from single particle analysis. After obtaining the sub-tomogram averages of C1 and C2, we fitted high-resolution C1 and C2 cryo-EM maps to the cryo-ET maps. According to the alignment information, the fitted maps were rotated and placed back to the original positions, resulting in the high-resolution maps of the entire S-shaped CA.

## Data availability

Cryo-EM maps and atomic coordinates have been deposited in the Electron Microscopy Data Bank under accession codes EMD-***** EMD-***** EMD-***** and EMD-***** and in the Protein Data Bank under accession codes **** and ****

## Acknowledgements

We thank S. Wu, K. Zhou, M. Llaguno at Yale University and K. Li and W. Huang at Case Western Reserve University for technical support on microscopy, J. Howard, G. Whitman and S. King for valuable discussion. This work was supported by start-up funds from Yale University, the National Institutes of Health grant R35GM142959 awarded to K.Z. and grant S10OD023603 awarded to F. Sigworth, and Rudolf J. Anderson Fellowship awards to L.H., and Y.W.

## Author contributions

L.H. prepared central apparatus samples; L.H., K.Z., P.C., R.Y., Y.W., and Q. R. collected cryo-EM and tomography data, determined the structures, built and refined the atomic models, and analyzed the structures; K.Z. wrote the paper with the help from L.H., Y.X. and all other co-authors.

## Supplementary Figures

**Supplementary Fig.1.**
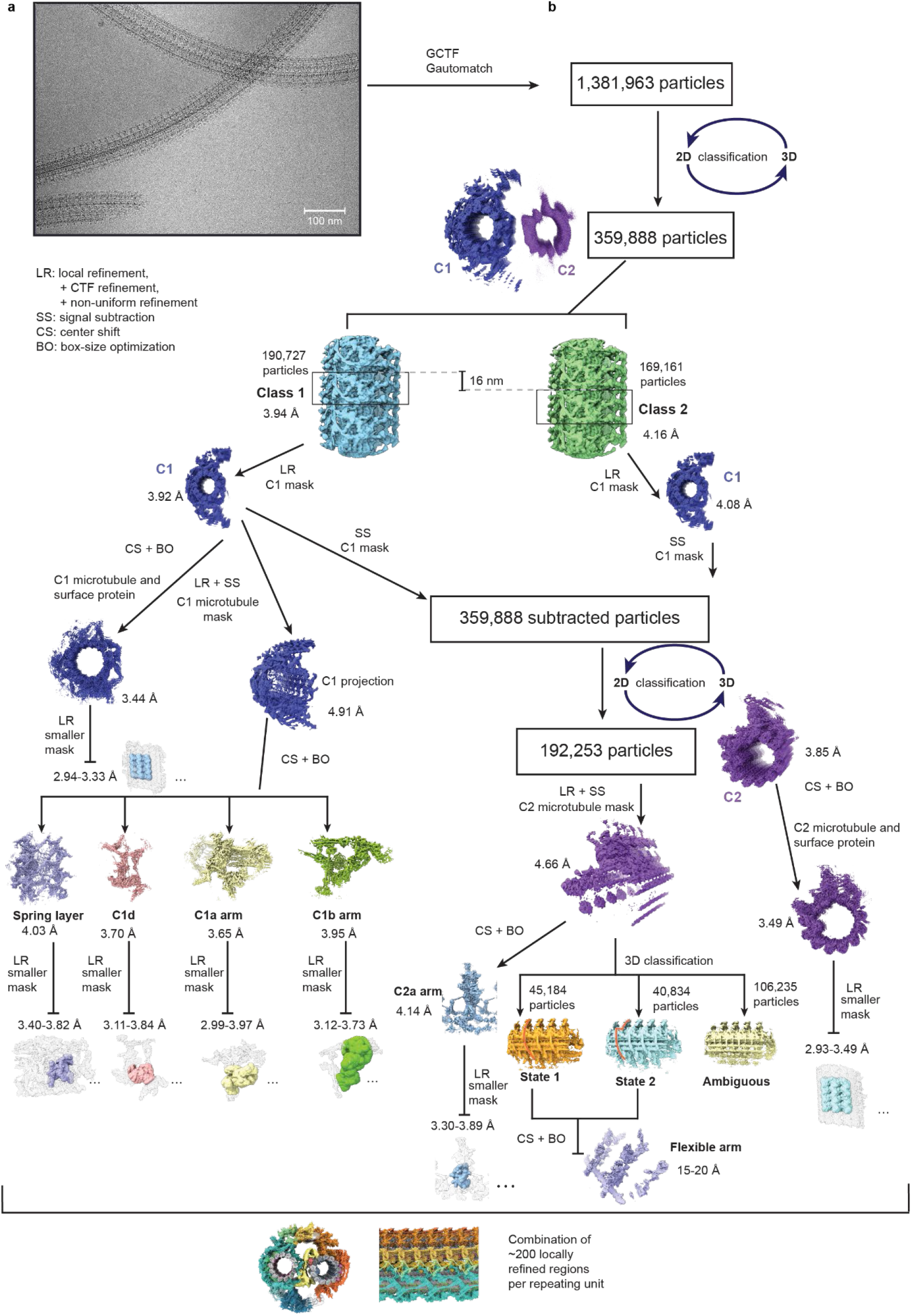
Data collection of CA and simplified workflow of data processing. (a) A representative image of isolated native CA from *C. reinhardtii*. (b) A simplified flowchart of cryo-EM data processing. ~10 rounds of 2D classification and 3D classification were performed iteratively to select high-quality particles for subsequent processing. As CA is an exceedingly large complex with extremely high flexibility, we utilized a multi-level refinement strategy that involved iterative local refinement (LR), signal subtraction (SS), center shift (CS), and box-size optimization (BO) to ‘divide and conquer’ this large structure with multiscale dynamics. The final composite map is combined from ~200 locally refined maps. A small subset of representative masks and density maps are shown here due to limited space. All intermediate results are available upon request.

**Supplementary Fig. 2.**
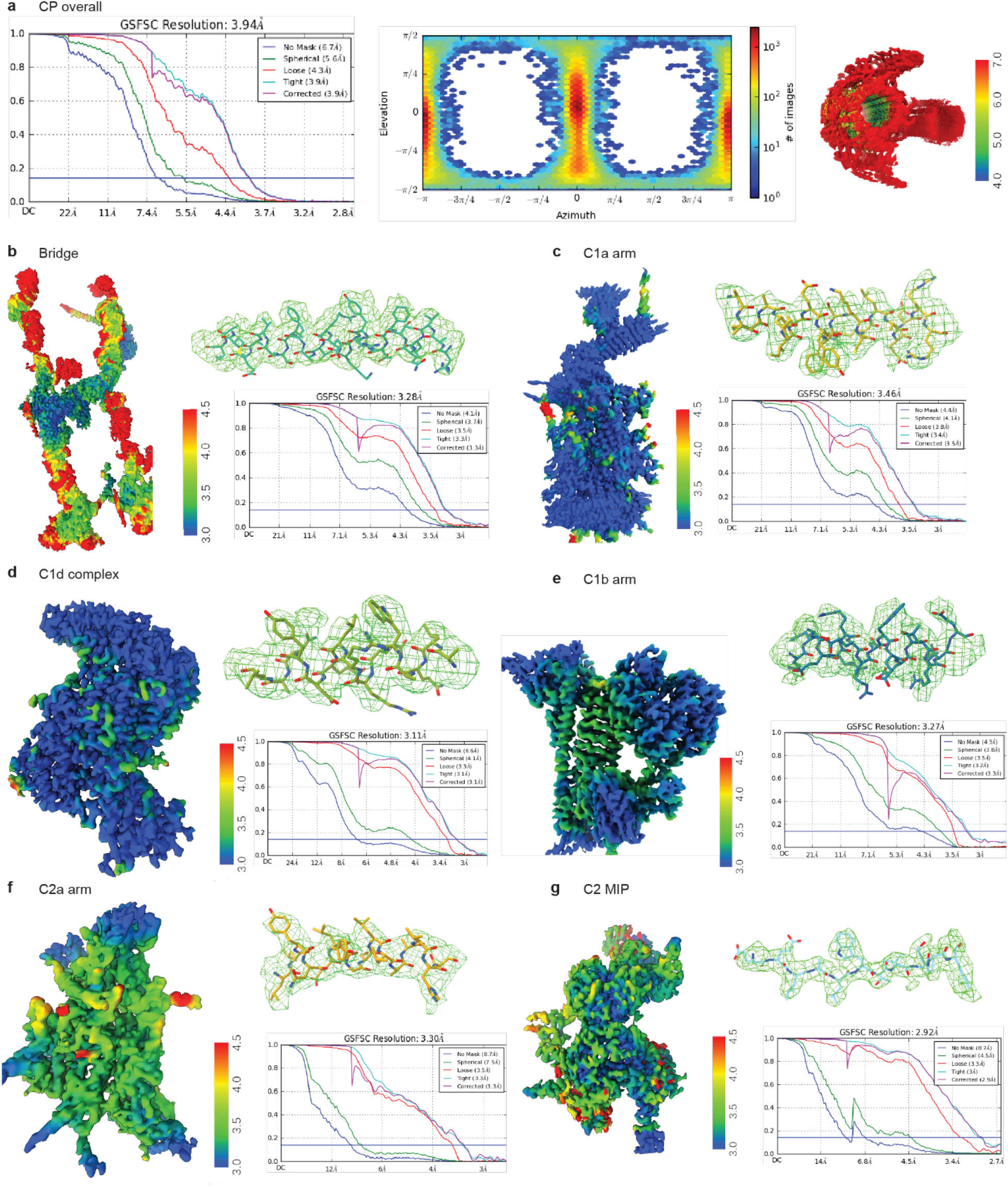
Evaluation of representative locally refined cryo-EM maps. (a) Overall evaluation of the globally refined CA structure. Fourier shell correction (FSC) curve (left) was estimated using a large mask that covers both halves. The angular distribution (middle) indicated that CA adopts preferred orientations, but sufficient for a 3D reconstruction. The effect was minimized after 2D and 3D classifications. Right: Local-resolution map of the reconstructed map of the CA after global refinement. As C1 dominates the alignment, the map is of C2 is completely blurred after global refinement due to continuous sliding between the two halves. (b-g) Representative local refinement results in different regions. Each panel contains a local-resolution map, the FSC curve, and a representative local density map with the atomic model built for different regions, including the bridge (b), C1a arm (c), C1d complex (d), C1b arm (e), C2a arm (f), and C2 MIP (g).

**Supplementary Fig. 3.**
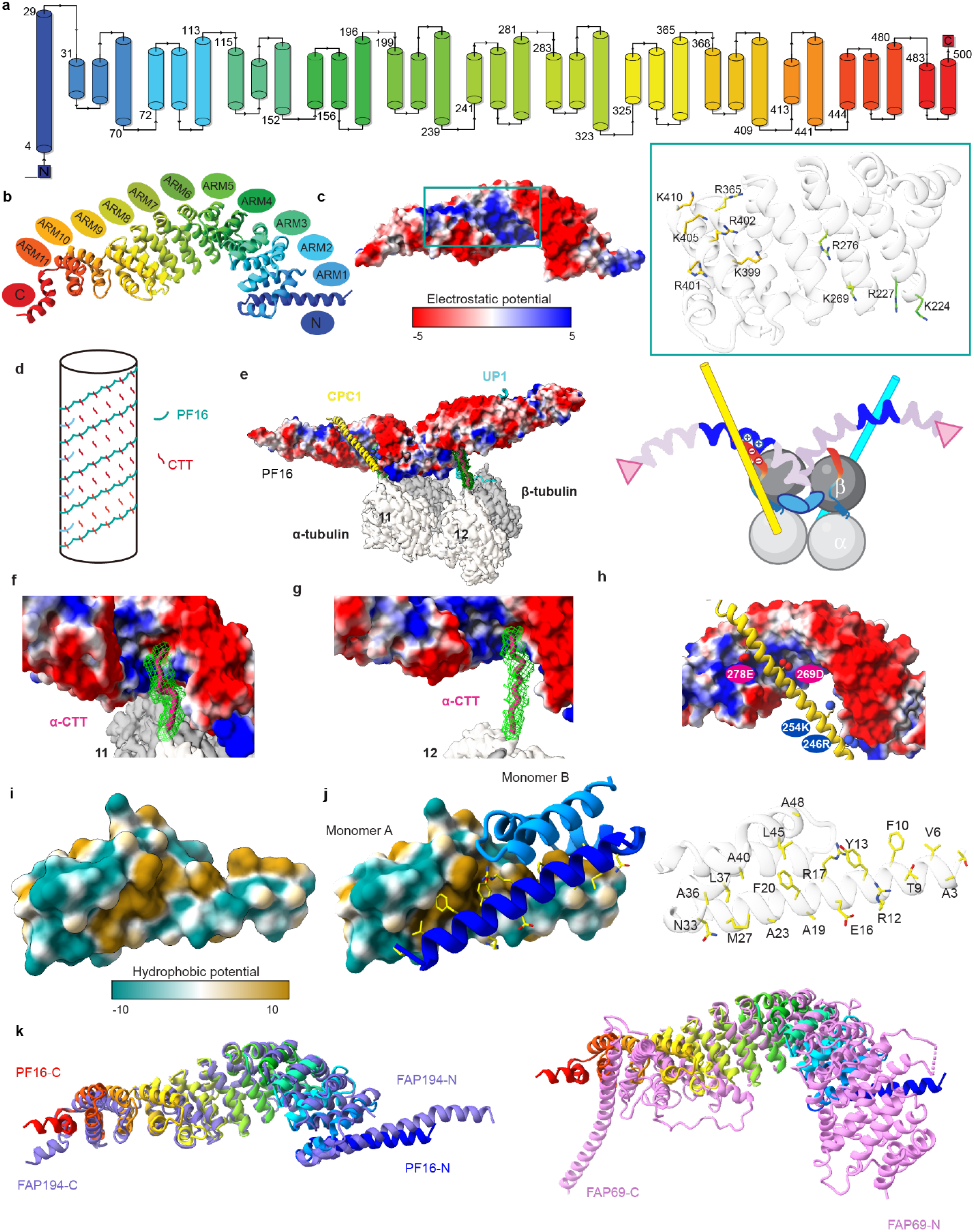
Characterization of the main armadillo protein PF16 structure. (a) A diagram of the PF16 topology. PF16 is mainly composed of α-helices, including 11 armadillo (ARM)-repeat motifs, a long N-terminal helix (residues 4-29), and two short C-terminal helices (483-489, 491-500). Each armadillo-repeat motif is ~40 amino-acid long, consisting of one long and two short helices except the 10^th^ repeat which is 29 amino-acid long containing a short helix and a long helix. (b) The atomic model of PF16. The PF16 structure contains an N-terminal helix (blue), 11 ARM motifs (ARM-1 to 11), and C-terminal helices (red). The domains are colored in rainbow from the N-terminus to the C-terminus. (c) The surface electrostatic potential map of PF16 (blue: positive, red: negative). The N-terminal helix and groove are positively charged, while the outer surface is negative charged. The blue box includes the details of positively charged residues, including 5 lysines (K) and 5 arginines (R). (d) A diagram shows how PF16 molecules wrap around the microtubule via their interaction with tubulin C-terminal tails (CTT). (e) A representative PF16 dimer bound to the C1 microtubule protofilament 11. A cartoon model is illustrated on the right to show the interactions. (f-g) Detailed views of the interactions between α-tubulin CTTs and positively charged grooves of PF16 dimer. (h) An enlarged view of the positively charged groove of PF16 that harbors CPC1. (i) Surface hydrophobic potential map of PF16 at the N-terminal region. (j) The dimerization interface of PF16. The surface hydrophobic potential map represents one subunit of the PF16 dimer while the cartoon represents the other. Dimerization of PF16 is mainly mediated by the hydrophobic cores of the two monomers, assisted by charged residues on the outer surfaces. The right panel shows the residues that contribute to the dimerization. (k) Superimposition of PF16 with the other two armadillo proteins, FAP194 (left) and FAP69 (right). PF16 is colored in rainbow, while FAP194 and FAP69 are colored in blue and violet, respectively. The structure of FAP194 is highly similar to that of PF16 and also contains 11 ARM repeats, one long N-terminal helix, and two short C-terminal helices. FAP69 is the largest among them, which has 13 armadillo-repeats followed by a helical domain (four short helices) and a long C-terminal helix.

**Supplementary Fig. 4.**
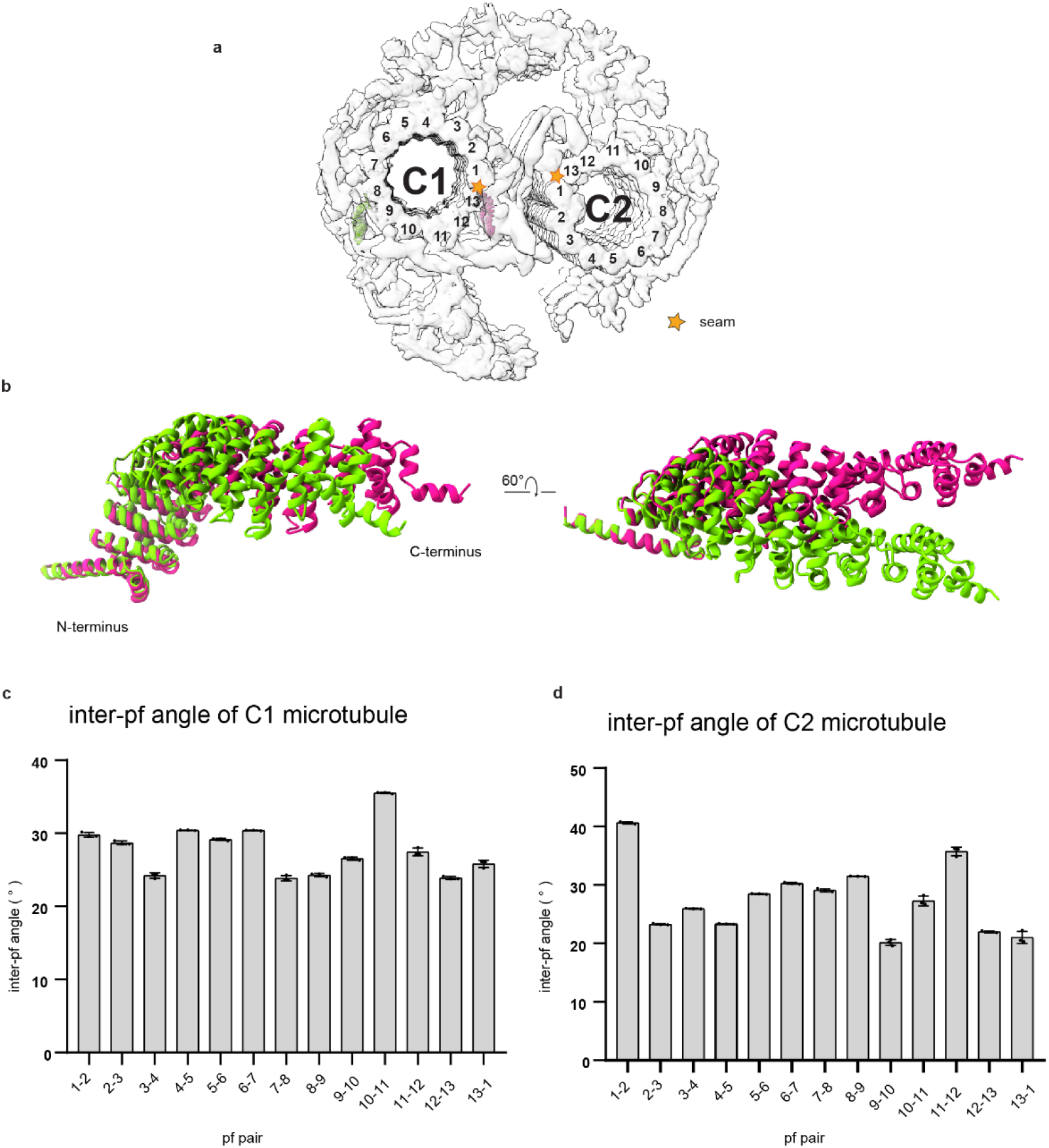
PF16 conformational changes and the inter-protofilament angles of the CA microtubules. (a) Cross-sectional view of CA showing locations of two PF16 monomers and seams. The protofilament (pf) number is labeled. (b) The conformational changes of PF16. A superposition of two PF16 monomers indicates they adopt different conformations without breaking the secondary structure. (The green molecule binds to pf8 and locates at C1d, while the pink molecule binds to pf13 and locates at bridge near the C1 seam.) (c) Plot of inter-pf angle of C1 microtubules. (d) Plot of inter-pf angle of C2 microtubules. Error bars represent Mean +/- SD.

**Supplementary Fig. 5.**
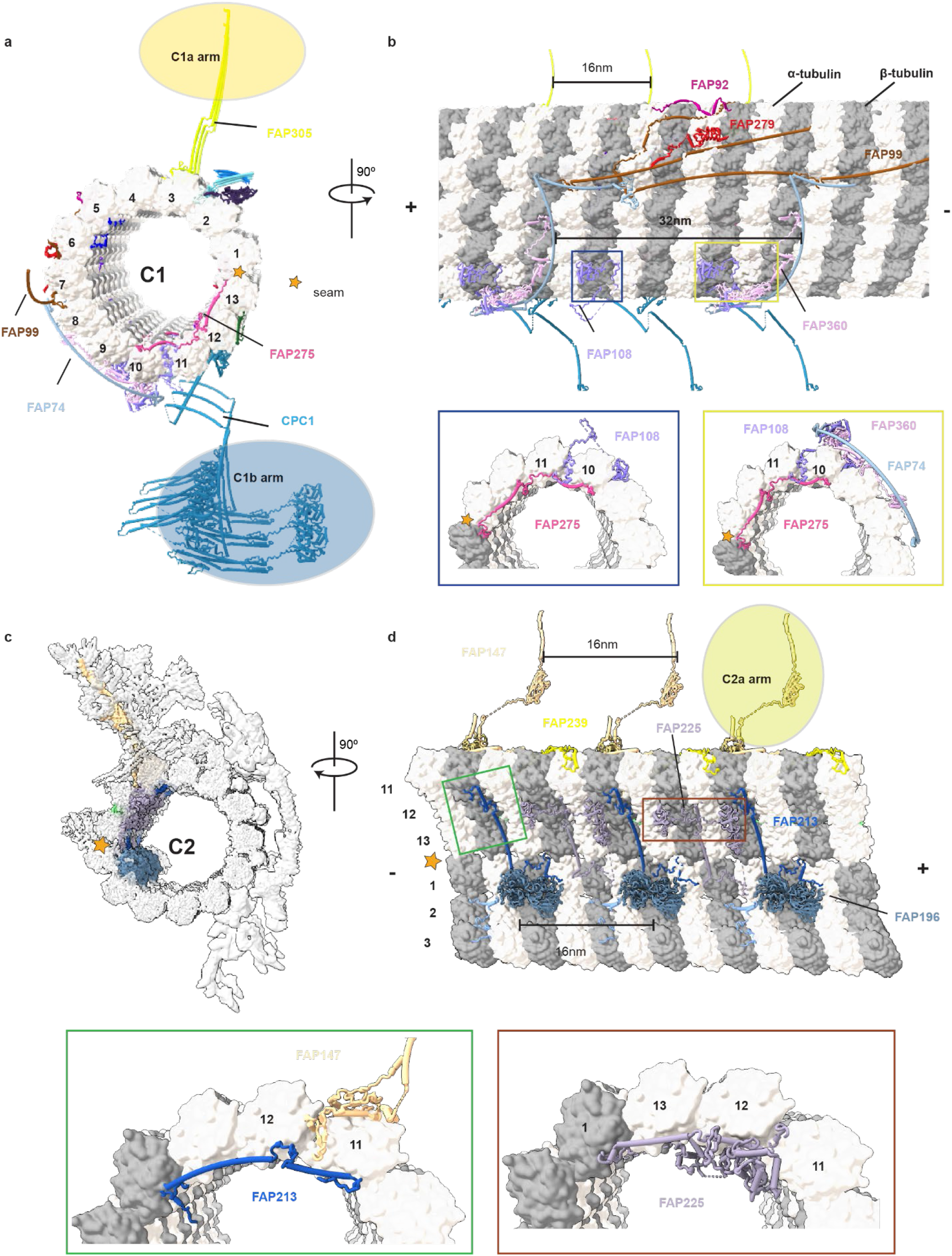
Microtubule surface proteins (MSPs) form interconnected networks around the CA microtubules. (a) Cross-sectional view of the C1 MSP network. The C1 microtubule is in surface representation. The rachis proteins (FAP305 and CPC1) indicate the positions of the C1a arm and C1b arm. The yellow star indicates the seam of the C1 microtubule. (b) Longitudinal view of the C1 outer surface proteins. The outer surface filament proteins (FAP74, FAP92, FAP99), FAP279, and FAP360 repeat at every 32 nm, while the EF-hand-like protein (FAP108) and rachis proteins (CPC1, FAP305) repeat at every 16 nm. The blue and yellow boxes indicate the site where FAP108 goes through the C1 microtubule wall to interact with the C1 MIPs. Blue box: the FAP108 penetrates through the cleft of protofilaments 10-11 and interacts with FAP275, the C1 MIP, which laterally lays on the C1 microtubule inner surface. Yellow box, FAP108 interacts with FAP275 in the lumen of the microtubule and tethers FAP74 and FAP360 on the outer surface. (c) Cross-sectional view of the C2 MSP network. (d) Longitudinal view of the C2 MSP. The C2 MSPs show a 16-nm periodicity. The rachis protein FAP147 indicates the position of C2a arm. FAP239 is a C2 MOSP that inserts a helix and loop into the cleft between protofilaments 10 and 11. The identified C2 MIPs include FAP196 (two WD40 domains), FAP225 (three EF-hand pairs), and FAP213 (long helices). The two sites where C2 MIPs and MOSPs are connected are indicated by green and brown boxes. Green box: FAP147 penetrates through the cleft between C2 microtubule protofilaments 11 and 12 to interact with FAP213, which interconnects the C2 MIPs and C2a arm. Brown box: the C2 MIP FAP225 protrudes out of the cleft to interact with C2 MOSPs.

**Supplementary Fig. 6.**
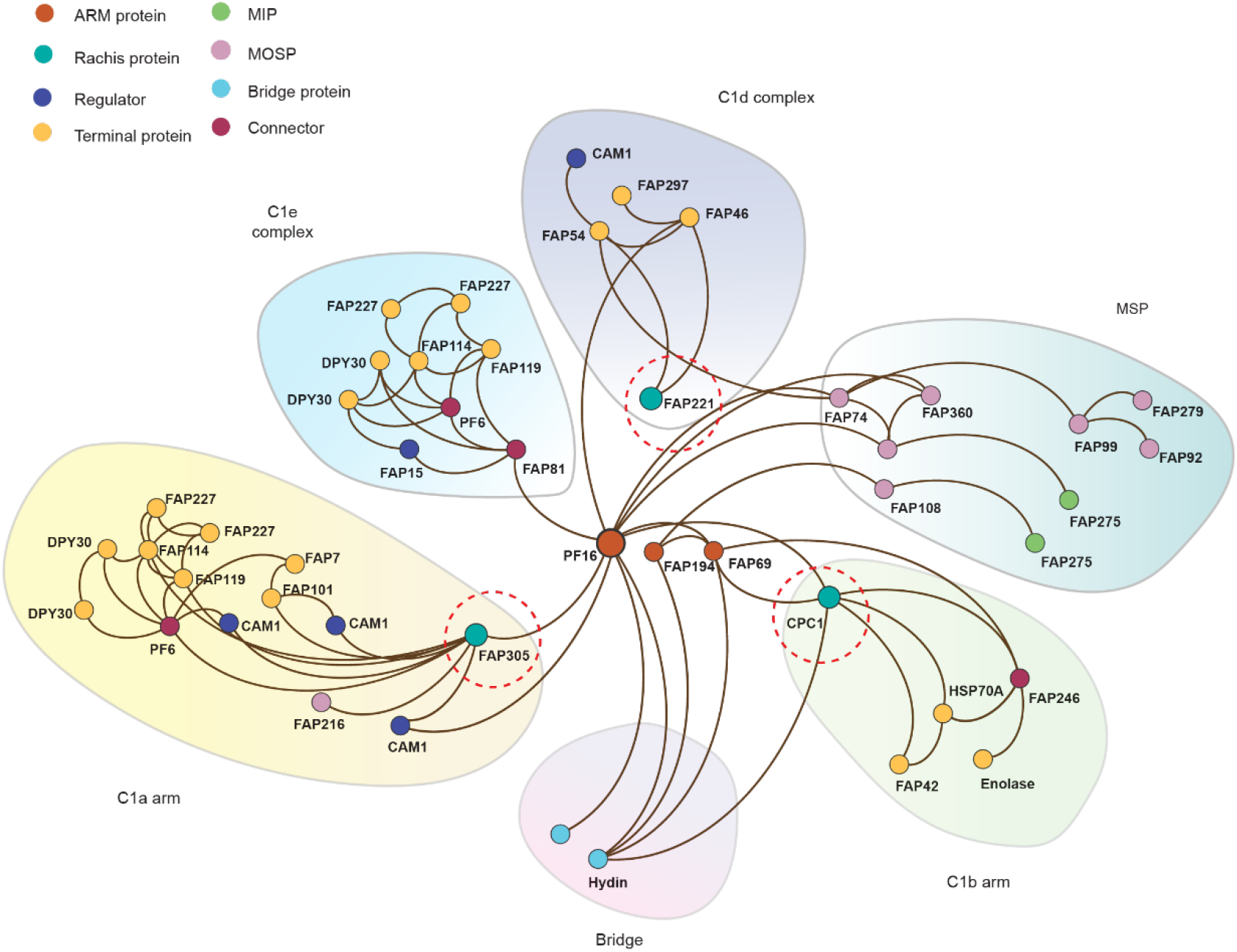
The interaction network of non-tubulin proteins on C1. A network shows the interactions of non-tubulin proteins associated with the C1 microtubule. The proteins are clustered into seven groups. The network is centered on the spring scaffold formed by armadillo-repeat proteins (PF16, FAP194, FAP69). To simplify the representation of the network, only one copy of each scaffold protein is displayed here. The six groups that are connected to the spring scaffold are colored differently. Proteins are colored differently based on their positions and functions. The rachis proteins (FAP305 in C1a arm; FAP221 in C1d complex; CPC1 in C1b arm) of the projections are colored in turquoise and encircled in red dashed lines.

**Supplementary Fig. 7.**
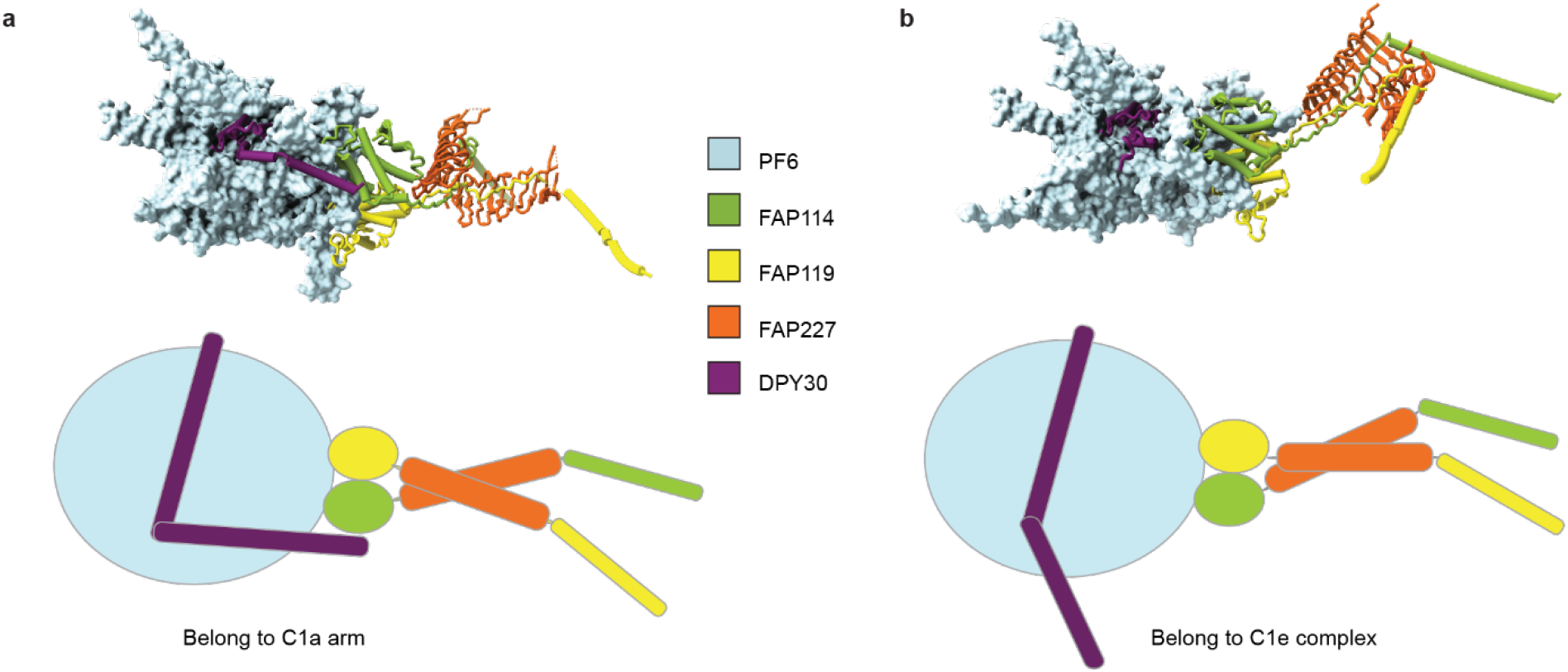
PF6 complex. Two copies of the PF6 complex in C1a and C1e regions. The two PF6 complexes consist of the same proteins but adopt different conformations. In C1a arm, the DPY30 dimer tightly binds to PF6, whereas one of the DPY30 subunits loosely binds to PF6 in C1e. The antennas-like subcomplexes, FAP114-FAP119-(FAP227)2, show different orientations in C1a arm (a) and C1e complex (b).

**Supplementary Fig. 8.**
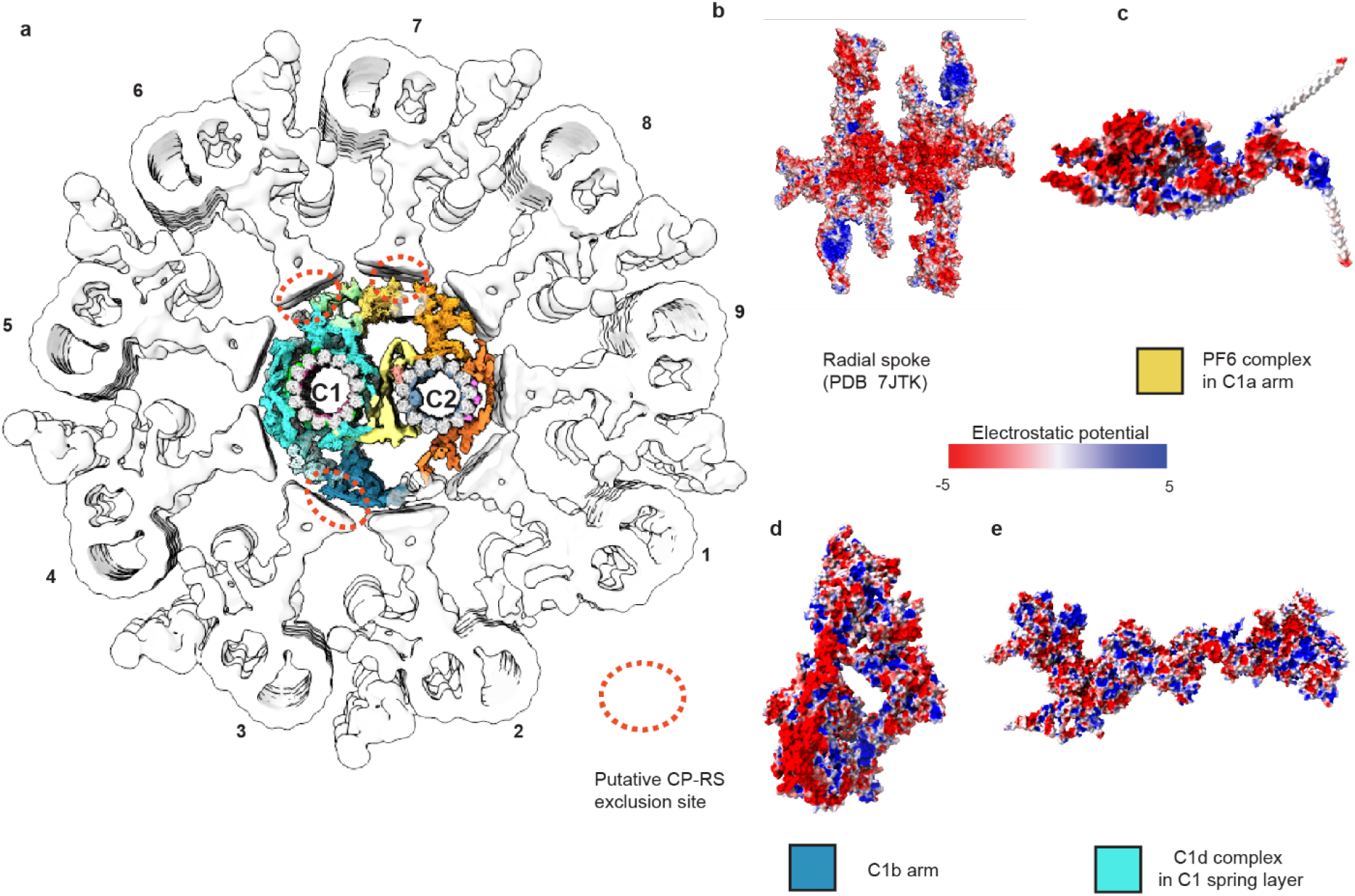
The architecture of the *C.reinhardtii* axoneme and the putative interacting regions between CA and radial spokes. (a) The architecture of *C.reinhardtii* axoneme. MTD-1 with associated structures (EMD: 2113), MTD2-8 with associated structures (EMD: 2132), and MTD-9 with associated structures (EMD: 2118) are used to composite the nine MTD and associated structures^15^. The red dashed ovals highlight potential repulsions between radial spokes and CA based on surface electrostatic potential analysis. (b-e) The major proteins involved in the repulsion interaction are rendered by surface electrostatic potential, including radial spoke head (b) (PDB: 7JTK)^94^, the PF6 complex in the C1a arm (c), C1b arm (d), and the C1d complex (e). The views show the CA projection surfaces that face radial spokes for possible interactions.

**Supplementary Fig. 9.**
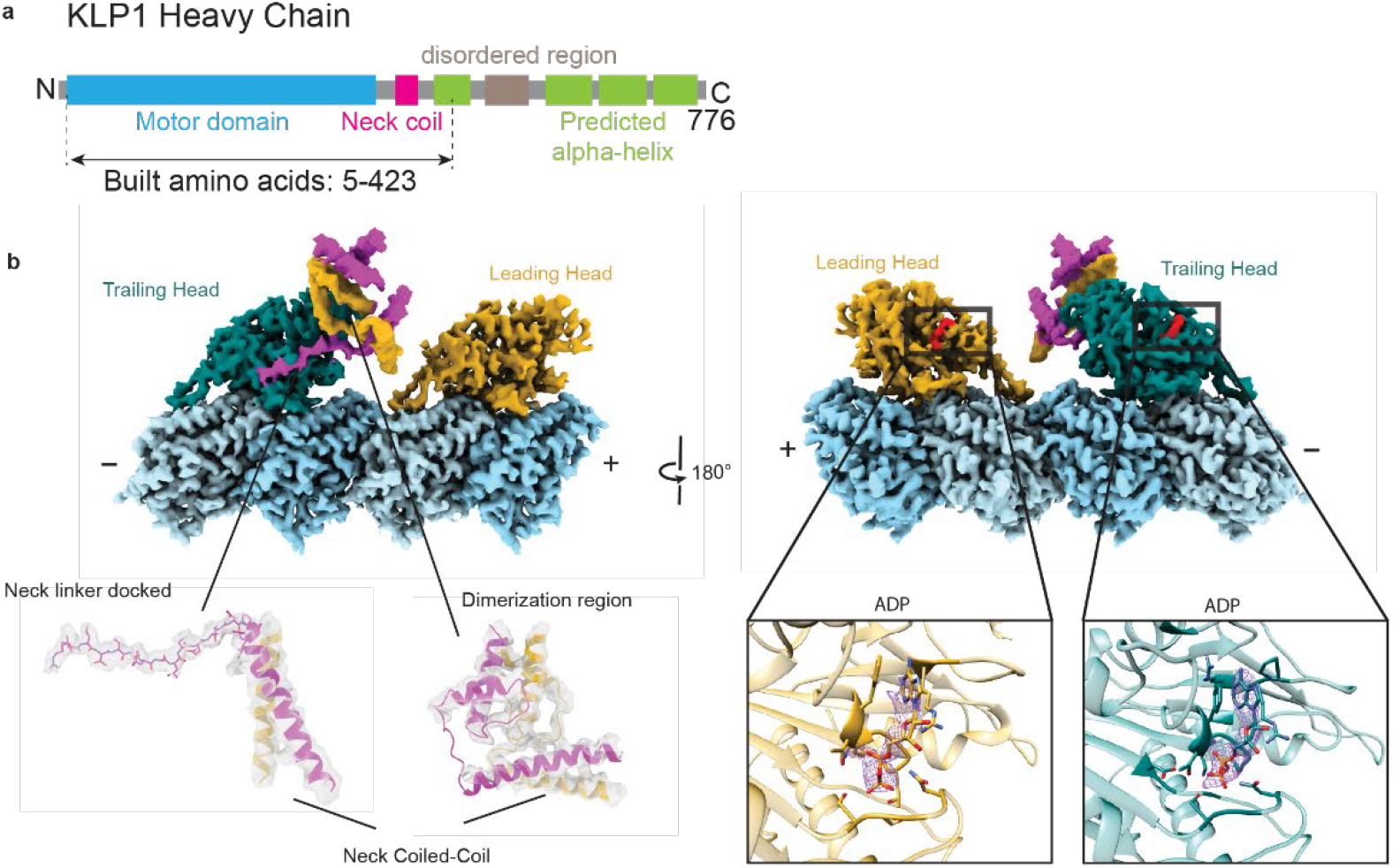
The structure of KLP1. (a) Domain organization of KLP1. KLP1 heavy chain contains a motor domain, a coiled-coil neck domain, and a tail region of three long alpha-helices. The tail and the neck are connected by a highly flexible region without ordered structures. We built the atomic models for the motor domain and neck domain and tentatively assigned the tail region. The near-atomic structures unambiguously show that two KLP1 heads adopt different conformations. (b) Two opposite views of the heads of the KLP1 dimer. The left panel shows that the trailing head and the leading head are dimerized by the neck coiled-coil, while the neck-linker of the trailing head is docked onto the motor head. The right panel shows that both heads bind ADP molecules at their nucleotide-binding sites.

**Supplementary Fig. 10.**
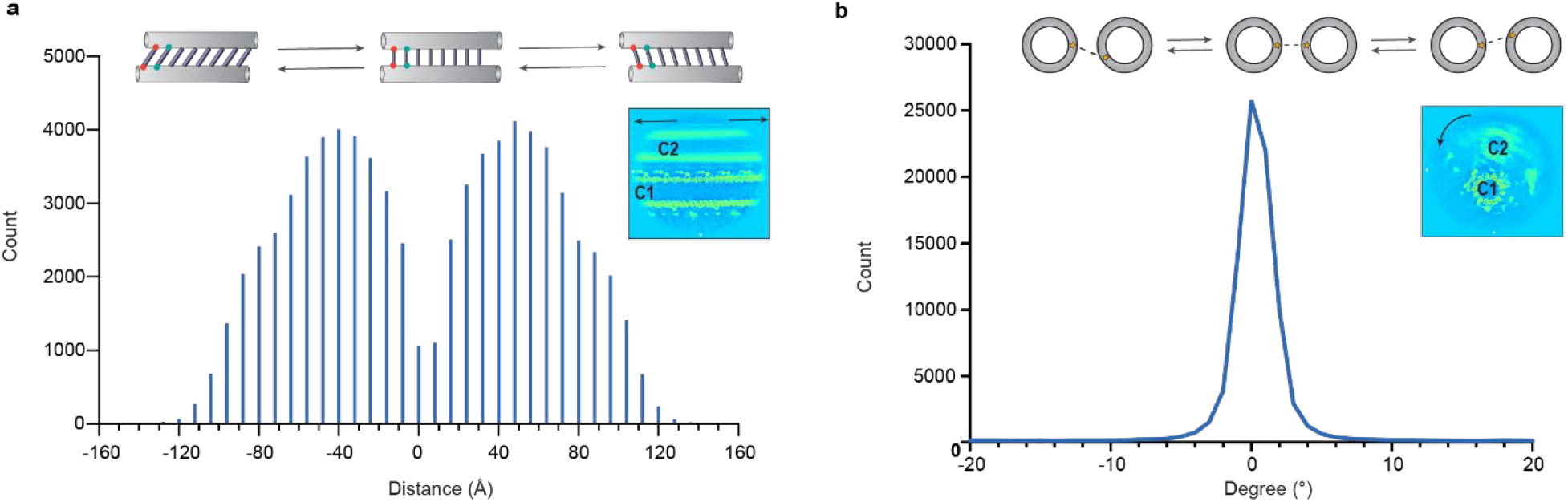
Relative movement between the two halves. (a) Statistics of the relative sliding between the two CA halves. A cartoon model and a slice of the cryo-EM map are illustrated to show the relative sliding. The longitudinal shift distance between the two microtubules distributes in a bell-like curve with a peak shift distance at ~4 nm and a maximum shift up to ~12 nm. (b) Statistics of the relative rotation between C1 and C2 microtubules. The standard deviation of the relative rotation angles is around ±2 degrees.

**Supplementary Fig. 11.**
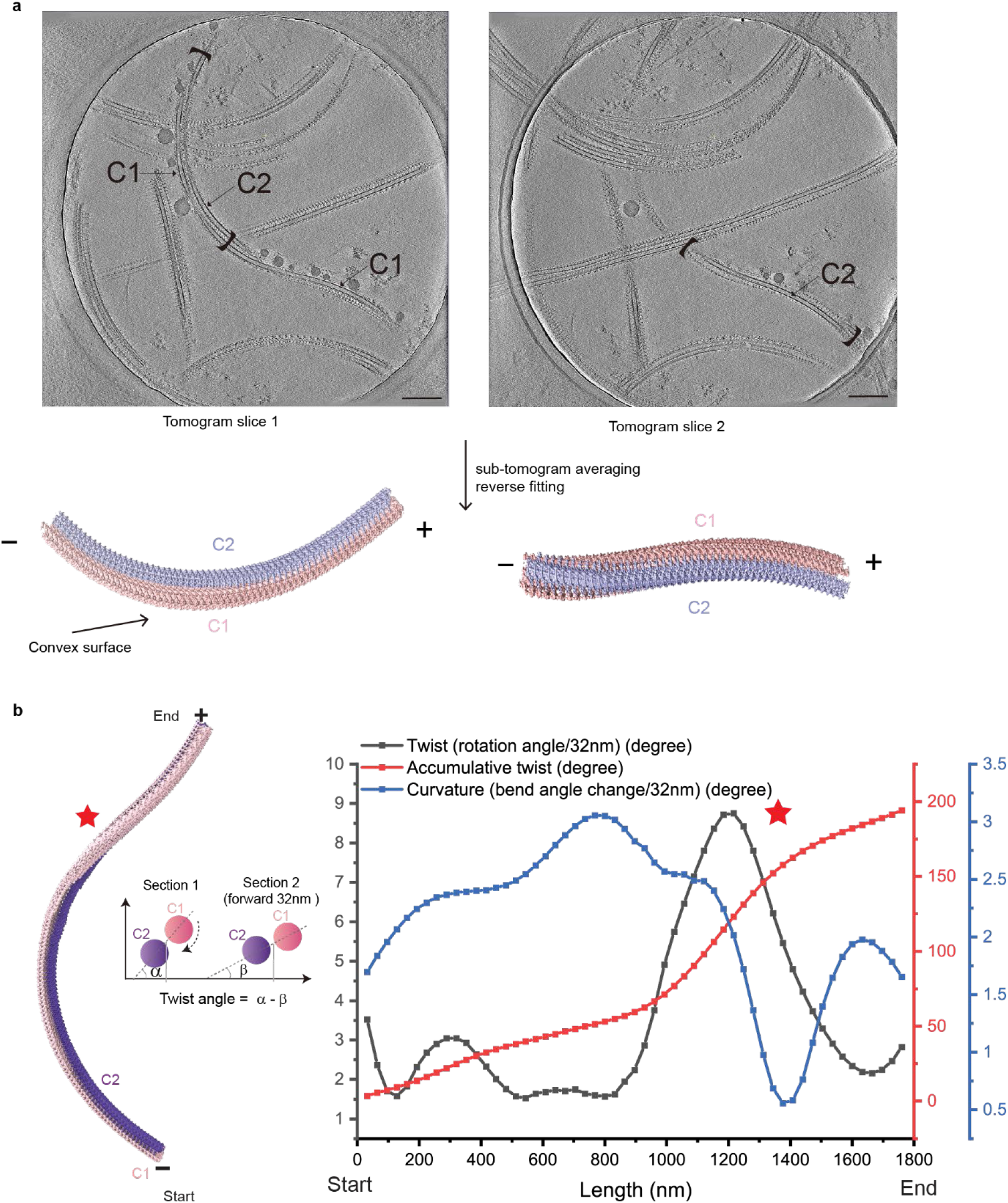
Cryo-electron tomography analysis of isolated CA. (a) Tomogram slices showing an ‘S-curved’ CA. Sub-tomogram averaging reveals that the C1 microtubule is always near the outer edge. In addition, a clockwise rotation (viewed from minus-end to plus-end) between C1 and C2 microtubules always exists throughout the entire CA. (b) Quantification of the CA geometry at a large scale. Black curve: relative twist angles between C1 and C2 per 32 nm indicate the local sharpness of the CA rotation. Red curve: accumulative twist angles of CA give the total amount of rotation from the start. Blue curve: curvature of CA shows the bending state of the entire CA. The quantification suggests that the sharpest rotation between C1 and C2 occurs when CA is least bent (red star).

**Supplementary Fig. 12.**
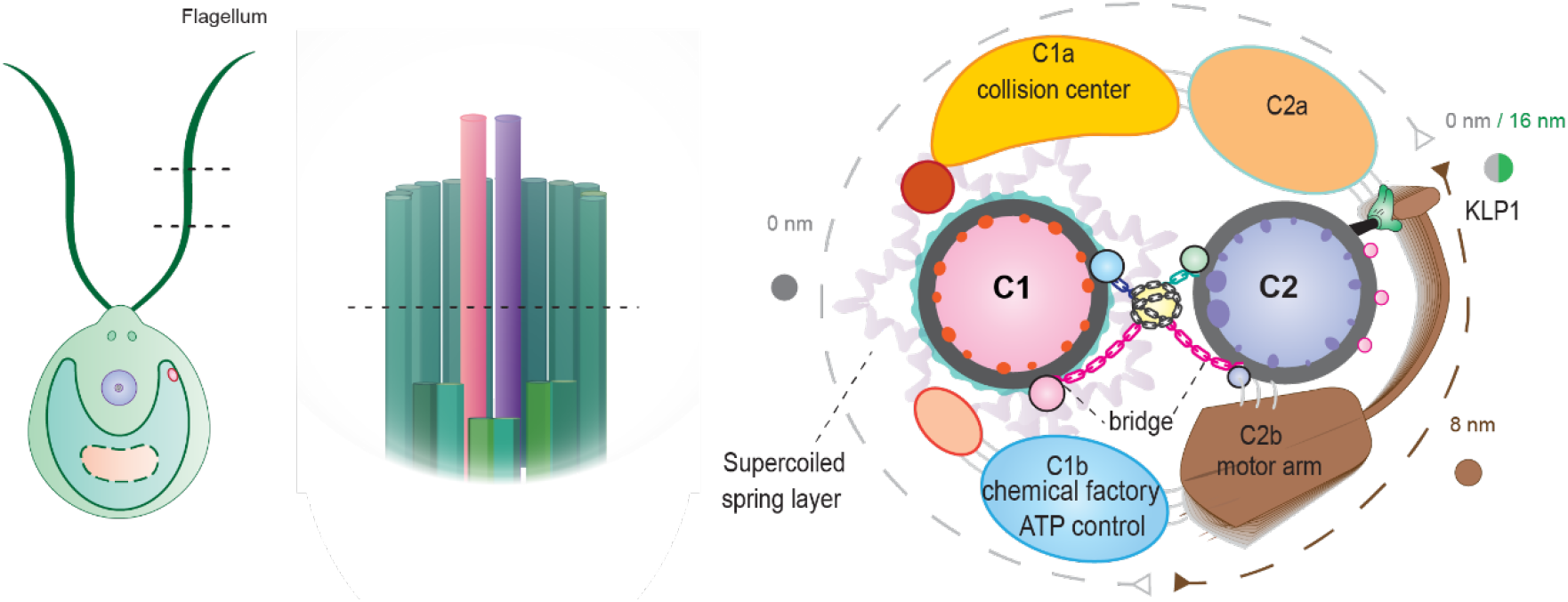
A proposed model CA. Schematics of the *C. reinhardtii* flagella (left), axoneme (middle), and CA architecture. The kinesin KLP1 is depicted as a ‘green hand’ to power the movement of the motor arm (brown oar), which likely plays a critical role in controlling CA conformations.

**Supplementary Fig. 13.**
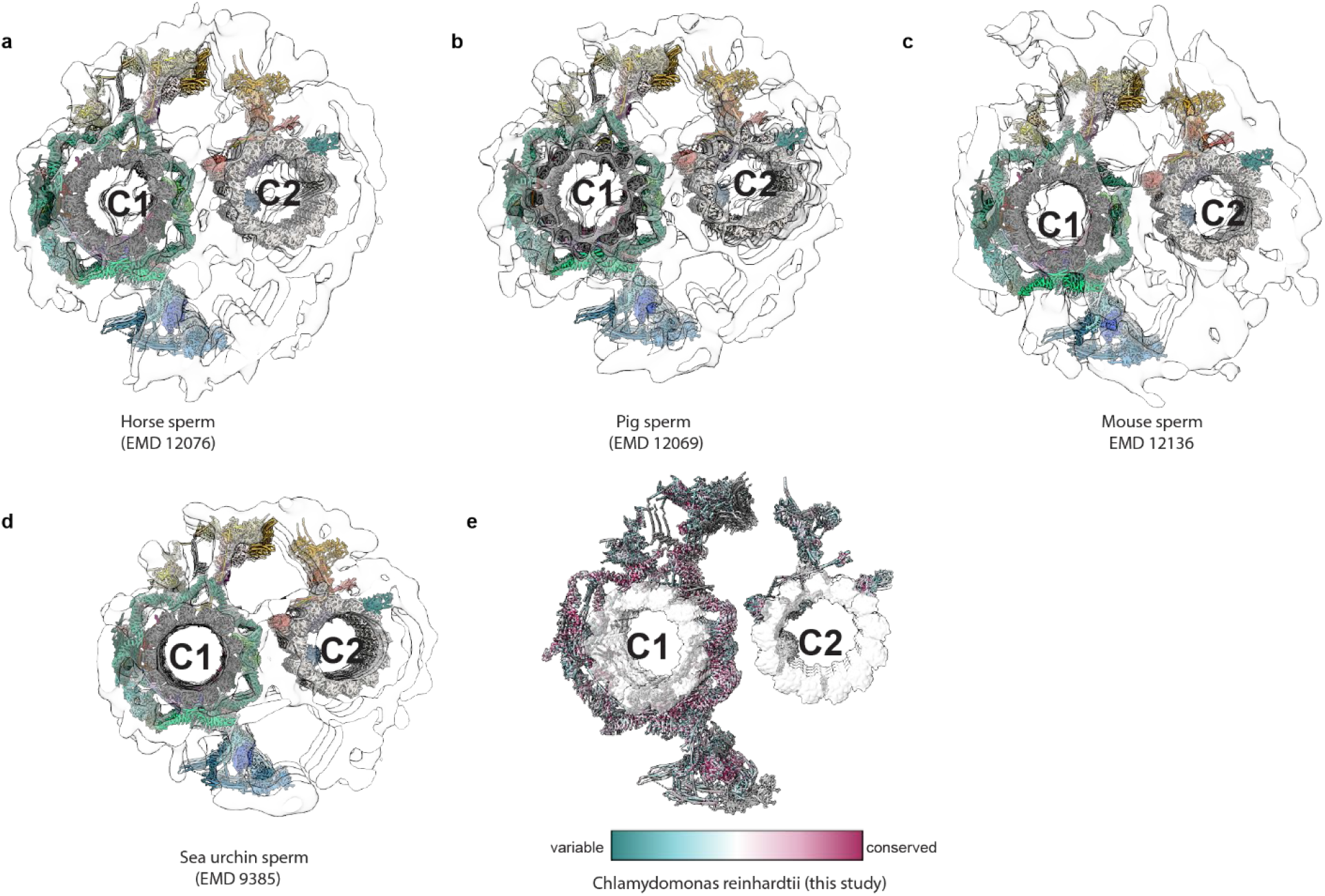
A comparison of CA structures among different species. (a-d) Cross-sectional views of the 32-nm repeat of CAs with atomic models docked into the cryo-ET maps. The docked models show that the core structures of CA are conserved across species, despite variations of the peripheral projections. The compared structures include (a) horse sperm (EMD-12076)^75^, (b) pig sperm (EMD-12069)^75^, (c) mouse sperm (EMD-12136)^75^, and (d) sea urchin sperm (EMD-9385)^30^. The color scheme of the atomic model of *C. reinhardtii* CA follows that of Fig1. (e) A conservation map of non-tubulin proteins of CA projections across species. Quantification of the conservation is calculated by sequence alignment of CA projection proteins from *C. reinhardtii* (taxid:3055), *Leishmania* (taxid:5658), *Mus musculus* (taxid:10090), *Danio rerio* (taxid:7955), *Silurana (Xenopus) tropicalis* (taxid:8364), *Drosophila melangaster* (taxid:7227), *Pan troglodytes* (taxid:9598), *Canis lupus* (taxid:9612), *Bos taurus* (taxid:9913), *Macaca mulatta* (taxid:9544), *Rattus norvegicus* (taxid:10116), *Gallus gallus* (taxid:9031), and *Homo sapiens* (taxid:9606) in UCSF ChimeraX^92^.

**Supplementary Table 1: Cryo-EM data collection, refinement, and validation statistics.**

**Supplementary Table 2: Identified proteins of the C. reinhardtii Central Apparatus.**

**Supplementary Video 1: PF16 conformational changes between the subunit at C1d and bridge regions.**

The movie shows that the PF16 molecules can elastically change its conformations without breaking the secondary structures. This conformational change is a reflection of its ability to adapt to the rotation angles between adjacent microtubule protofilaments. It also indirectly suggests the potential of PF16 molecule to undergo an elastic conformational change upon local curvature changes during ciliary beating.

**Supplementary Video 2: Continuous sliding between the two halves of CA.**

The movie was generated by aligning all repeats from the same CA. The pink and blue line segments are both 80 nm long. The starts and ends of these line segments in all frames represent the centers of equivalent structural features of the repeating units. The movie shows that the relative sliding between the two halves is accumulated along CA in the curved region.

**Supplementary Material 1: Structural comparison between CA proteins and their human orthologs.**

**Table 1:** Cryo-EM data collection, refinement and validation statistics.

| **Data collection and processing** | Dataset 1 | Dataset 2 | Dataset 3 |
| --- | --- | --- | --- |
| Microscopy | Titan Krios G2 | Titan Krios G2 | Titan Krios G3i |
| Detector | K2 Quantm (Gatan) | K2 Quantm (Gatan) | K3 (Gatan) |
| Magnification | 130,000 | 105,000 | 65,000 |
| Voltage (kV) | 300 | 300 | 300 |
| Electron exposure (e-/Å<sup>2</sup>) | 39.2 | 48.4 | 38.6 |
| Defocus range (μm) | -1.2 to -2.5 | -1.2 to -2.5 | -1.2 to -2.5 |
| Pixel size (Å) | 1.05 | 1.33 | 1.33 |
| Symmetry imposed |  | C1 |  |
| Initial particle number |  | 1,381,963 * |  |
| Final particle number |  | 359,888 ** |  |
| Final particles number | 190,727 | 192,253 |  |
|  | #1 C1 projections and MIPs<br>(EMDB-24207)<br>(PDB-7N6G) | #2 C2 projections and MIPs<br>(EMD-24191)<br>(PDB-7N61) |  |
| Map resolution (Å) | 3.5 | 3.8 |  |
| FSC threshold | 0.143 | 0.143 |  |
| Map resolution range (Å) | 2.8 - 6 | 2.8 - 7 |  |
| **Refinement** |  |  |  |
| Initial model used (PDB code) | Ab initio | Ab initio |  |
| Model resolution (Å) |  |  |  |
| FSC threshold | 0.143 | 0.143 |  |
| Model resolution range (Å) | 3.0 - 4.0 | 3.0 - 5.0 |  |
| Map sharpening B factor (Å<sup>2</sup>) | -80 | -80 |  |
| Model composition | C1 projections, C1 microtubule<br>and its associated MIPs. | C2 projections, C2 microtubule and<br>its associated MIPs. |  |
| Non-hydrogen atoms |  |  |  |
| Protein residues | 793,760 | 444,292 |  |
| Ligands | 106,386 | 57,482 |  |
|  | GTP/GDP/Mg<sup>2+</sup> (52/52/52). | GTP/GDP/Mg<sup>2+</sup> (52/52/52), ADP (4) |  |
| B factors (Å<sup>2</sup>) |  |  |  |
| Protein | 103.11 | 98.66 |  |
| Ligand | 102.71 | 101.14 |  |
| R.m.s. deviations |  |  |  |
| Bond lengths (Å) | 0.0080 | 0.004 |  |
| Bond angles (°) | 0.977 | 0.617 |  |
| Validation |  |  |  |
| MolProbity score | 2.30 | 1.78 |  |
| Clashscore | 17.7 | 12.21 |  |
| Poor rotamers (%) | 1.57% | 0.06% |  |
| Ramachandran plot |  |  |  |
| Favored (%) | 95.38% | 96.97% |  |
| Allowed (%) | 4.19% | 3.02% |  |
| Disallowed (%) | 0.42% | 0.01% |  |
| Rama Z-score | -1.33±0.02 | 0.33±0.03 |  |

\* All three datasets were merged and processed together.
\*\* Final particle number before focused refinement.

**Supplementary Table 2.** Identified proteins of the *C. reinhardtii* Central Apparatus.

| Central Apparatus Proteins | Phytozomev5 gene or UniProt ID | Molecular mass in Da (number of residues) | Locations / Copies per repeating unit* (subunit periodicity) | Built residues | Side chain confidence (1 high, 2 moderate, 3 no) | local resolutions for identified regions (Å) |
| --- | --- | --- | --- | --- | --- | --- |
| **C1 associated proteins** (C1 repeating unit periodicity, 32 nm) |  |  |  |  |  |  |
| CAM1 | Cre03.g178150.t1.1 | 18,296 (163) | C1a / 6 (16 nm)<br>C1d / 1 (32nm) | C1a arm: 6-151<br>C1d: 4-152 | 1 | 3.5 |
| CPC1 | Cre03.g183200.t1.2 | 205,137 (1929) | C1b / 2 (16 nm) | 2-292, 322-505, 533-635, 683-719, 764-986, 994-1103, 1153-1341, 1356-1396, 1434-1676, 1741-1843, 1896-1929 | 1 | 3.0 (MT surface)<br>4.0 projection |
| DPY30 | Cre06.g279100.t1.2 | 11,483 (110) | C1a / 4 (16 nm)<br>C1e / 2 (32 nm) | C1a arm: 24-87<br>C1e: 36-90/7-90 | 1 | 3.0 |
| Enolase | Cre12.g513200.t1.2 | 51,602 (477) | C1b / 4 (16 nm) | 2-477 | 1 | 3.3 |
| FAP7 | Cre12.g531800.t1.1 | 54,654 (507) | C1a / 8 (16 nm) | 33-127, 187-225, 232-507 | 1 | 3.0 |
| FAP15 | Cre06.g292550.t1.2 | 34848 (304) | C1e / 1 (32 nm) | 2-296 | 1 | 3.5 |
| FAP42 | Cre12.g519950.t1.1 | 268,848 (2540) | C1b / 2 (16 nm) | 1235-1375, 1378-1723, 1790-1851, 1894-2215, 2231-2325, 2356-2374, 2395-2417, 2433-2499 | 2 | 3.6 |
| FAP46 | Cre10.g420800.t1.1 | 289,251 (2784) | C1d / 1 (32 nm) | 2-208, 225-270, 298-372, 375-620, 623-668, 701-744, 786-833, 888-910, 913-925, 933-952, 957-1017, 1024-1274, 1343-1583, 1695-1993, 1997-2073, 2098-2229, 2234-2275, 2305-2348, 2411-2442, 2465-2491, 2493-2507, 2531-2563, 2588-2734, 2756-2765 | 1 | 3.2 |
| FAP47 | Cre17.g704300.t1.1 | 319,659 (2339) | C1 bridge / 2 (16 nm) | 3-208, 220-261, 298-326, 344-369, 371-613 | 2 | 3.5 |
| FAP54 | Cre12.g518550.t1.1 | 289,251 (2784) | C1d / 1 (32 nm) | 5-290, 292-302, 317-387, 409-476, 497-536, 632-810, 817-935, 942-1108, 1121-1441, 1453-1535, 1587-1667, 1725-1780, 1821-1929, 1986-2323, 2350-2376, 2435-2769, 2804-2903, 2950-3150, 3182-3222 | 1 | 3.2 |
| FAP69 | Cre03.g168200.t1.2 | 114,244 (1102) | C1 armadillo scaffold 2 (16 nm) | 11-147, 161-197, 205-293, 491-714, 745-997, 1002-1064 | 2 | 3.6 |
| FAP74 | Cre06.g271150.t1.1 | 203,808 (1940) | C1 MOSP / 1 (32 nm) | 102-502 | 1 | 3.3 |
| FAP76 | Cre09.g387689.t1.1 | 172,467 (1638) | C1 armadillo scaffold / 1 (32 nm) | 101-158, 169-221, 226-326, 335-377, 399-489, 498-562, 564-685, 690-738, 745-1072, 1105-1166 | 3 | 8 |
| FAP81 | Cre06.g296850.t1.2 | 234,211 (2215) | C1e / 1 (32 nm) | 280-319, 322-347, 353-439, 445-584, 685-745, 760-973, 1048-1095, 1101-1118 | 1 | 3.5 |
| FAP92 | Cre13.g562250.t1.1 | 153,520 (1471) | C1 MOSP / 1 (32 nm) | 1135-1221 | 1 | 3.5 |
| FAP99 | Cre14.g624400.t1.2 | 90,188 (795) | C1 MOSP / 1 (32 nm) | 229-520, 525-645, 649-767 | 1 | 3.5 |
| FAP101 | Cre02.g112100.t1.1 | 85,717 (835) | C1a / 2 (16 nm) | 31-93, 154-182, 195-214, 220-556, 598-678, 692-744, 817-835 | 1 | 3.2 |
| FAP108 | Cre06.g297200.t1.2 | 47,727 (446) | C1 MOSP / 2 (16 nm) | 36-176, 178-209, 229-273, 277-304, 332-446 | 1 | 3.3 |
| FAP114 | Cre09.g389282.t1.1 | 31,531 (286) | C1a / 2 (16 nm)<br>C1e / 1 (32 nm) | 6-170, 175-225 | 2 | 3.2 |
| FAP119 | Cre06.g256450.t1.2 | 33,807 (306) | C1a / 2 (16 nm)<br>C1e / 1 (32 nm) | 2-171, 178-218 | 2 | 3.2 |
| FAP194 | Cre12.g522150.t1.1 | 57,706 (571) | C1 armadillo scaffold / 2 (32 nm) | 15-72, 77-522 | 2 | 3.5 |
| FAP216 | Cre12.g497200.t1.2 | 79,078 (739) | C1 MOSP / 2 (16 nm) | 520-580 | 1 | 3.4 |
| FAP221 | Cre11.g476376.t1.1 | 109,272 (1023) | C1d / 1 (32 nm) | 685-931, 976-1023 | 1 | 3.5 |
| FAP227 | Cre17.g729850.t1.2 | 19,525 (173) | C1a / 4 (16 nm)<br>C1e / 2 (32 nm) | 20-39, 70-173 | 2 | 4.0 |
| FAP246 | Cre14.g618750.t1.1 | 120,322 (1138) | C1b / 2 (16 nm) | 47-368, 378-417, 478-530, 560-933, 941-1055 | 1 | 3.5 |
| FAP275 | Cre05.g239200.t1.2 | 18,142 (168) | C1 MIP / 2 (16 nm) | 8-168 | 1 | 3.5 |
| FAP279 | Cre06.g268650.t1.1 | 43,207 (401) | C1 MOSP / 1 (32 nm) | 18-166, 172-219 | 1 | 3.8 |
| FAP289 | Cre01.g009800.t1.2 | 46,264 (447) | C1 MOSP / 1 (8 nm) | 118-222 | 1 | 3.5 |
| FAP297 | Cre01.g029350.t1.2 | 99,058 (945) | C1d / 1 (32 nm) | 359-489, 524-566, 596-631, 635-654, 657-787, 790-840, 846-870 | 1 | 3.9 |
| FAP305 (MOT17) | Cre11.g482300.t1.2 | 28,490 (244) | C1a / 2 (16 nm) | 2-217 | 1 | 3.3 |
| FAP360 | Cre49.g761347.t1.1 | 80,656 (749) | C1 MOSP / 1 (32 nm) | 21-274, 300-328, 337-420 | 1 | 3.5 |
| HSP70 | Cre08.g372100.t1.2 | 71,214 (651) | C1b / 2 (16 nm) | 5-256, 260-616 | 1 | 3.4 |
| Hydin | Cre01.g025400.t1.2 | 528,365 (4929) | C1b / 1 (16 nm) | 12-340, 361-427 | 2 | 3.5 |
| PF6 | Cre10.g434400.t1.2 | 237,483 (2301) | C1a / 2 (16 nm)<br>C1e / 1 (32 nm) | C1a arm: 398-434, 505-624, 663-700, 739-979, 1066-1106, 1128-1206, 1270-1404, 1429-1489, 1510-1534, 1605-1742, 1752-1801<br>C1e: 404-434, 507-627, 664-887, 916-938, 954-978, 1067-1206, 1272-1404, 1429-1489, 1510-1534, 1605-1657, 1675-1821 | 1 | 3.5 |
| PF16 | Cre09.g394251.t1.1 | 54,655 (512) | C1 armadillo scaffold / 20 (32 nm) | 3-501 | 2 | 3.1 |
| **C2 associated proteins** (repeating unit periodicity, 16 nm) |  |  |  |  |  |  |
| FAP65 | Cre07.g354551.t1.2 | 233,006 (2257) | C2a / 1 (16 nm) | 1108-1425, 1443-1537, 1561-1777 | 1 | 3.6 |
| FAP70 | Cre07.g345400.t1.2 | 111,003 (1074) | C2a / 2 (16 nm) | 397-447, 454-705, 751-1071 | 2 | 3.7 |
| FAP147 | Cre04.g224250.t1.1 | 102,004 (976) | C2a / 1 (16 nm) | 90-115, 129-366, 409-592 | 1 | 3.3 |
| FAP178 | Cre10.g418150.t1.2 | 23,482 (222) | C2 bridge / 2 (16 nm) | 27-154 | 1 | 3.3 |
| FAP196 | Cre17.g728650.t1.1 | 64,814 (618) | C2 MIP / 1 (16 nm) | 2-618 | 1 | 3.2 |
| FAP213 | Cre16.g690100.t1.2 | 22,226 (201) | C2 MIP / 1 (16 nm) | 9-40, 44-199 | 1 | 3.3 |
| FAP225 | Cre01.g051050.t1.2 | 81,320 (758) | C2 MIP / 1 (16 nm) | 38-309, 316-424, 607-727, 741-754 | 1 | 3.1 |
| FAP239 | Cre03.g145387.t1.1 | 57,646 (528) | C2 MOSP / 1 (16 nm) | 56-221 | 2 | 3.4 |
| KLP1 | Cre02.g073750.t1.2 | 82,997 (776) | Motor arm / 2 (16 nm) | 4-423 | 2 | 3.5 |
| PF20 | P93107 (UniProt) | 65,839 (606) | C2 bridge / 2 (16 nm) | 64-224, 292-606 | 1 | 3.5 |

\* 32 nm for C1 proteins and 16 nm for C2 proteins.

### Supplementary materials

Structural comparison between CA proteins and their human orthologs.

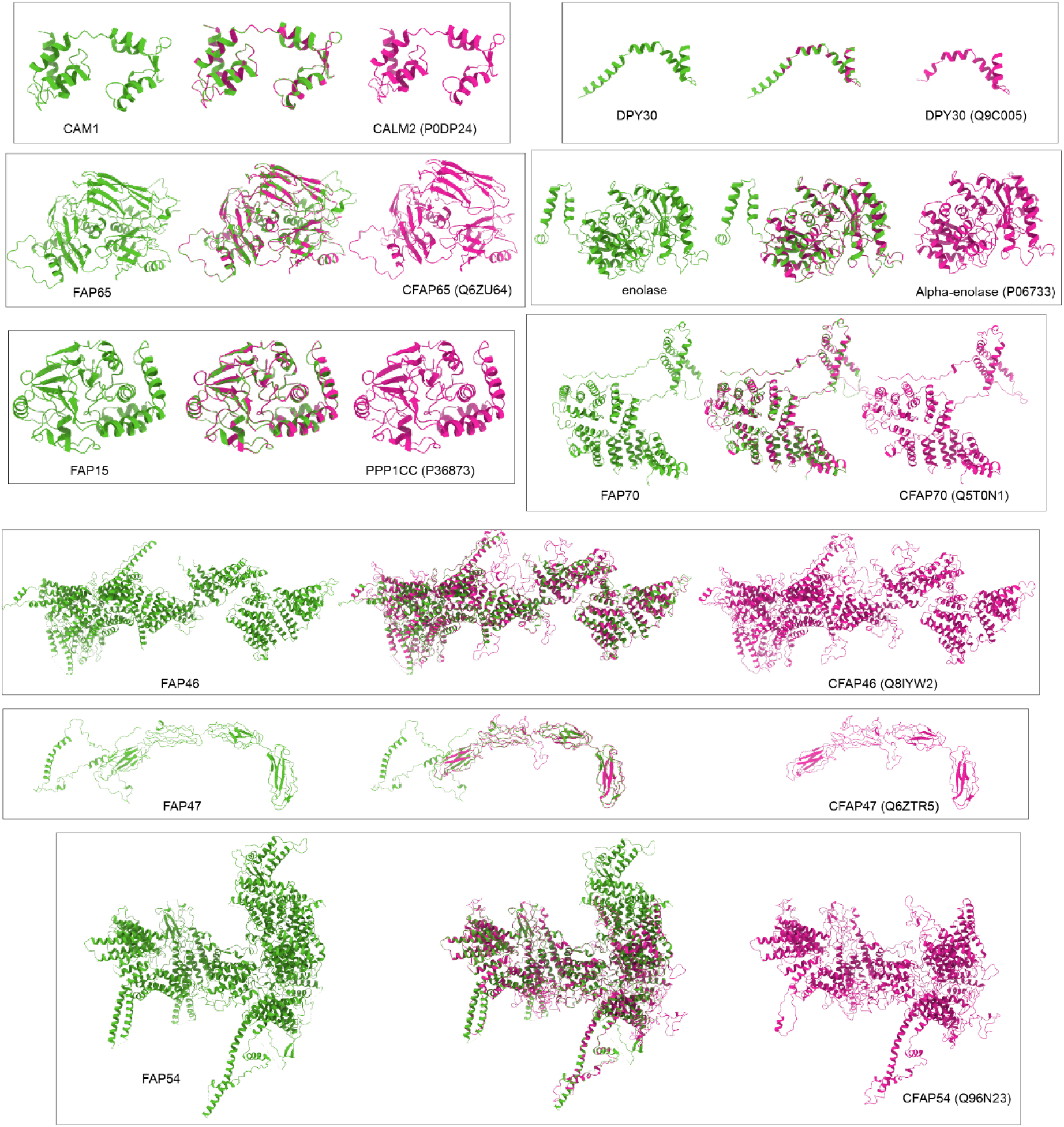
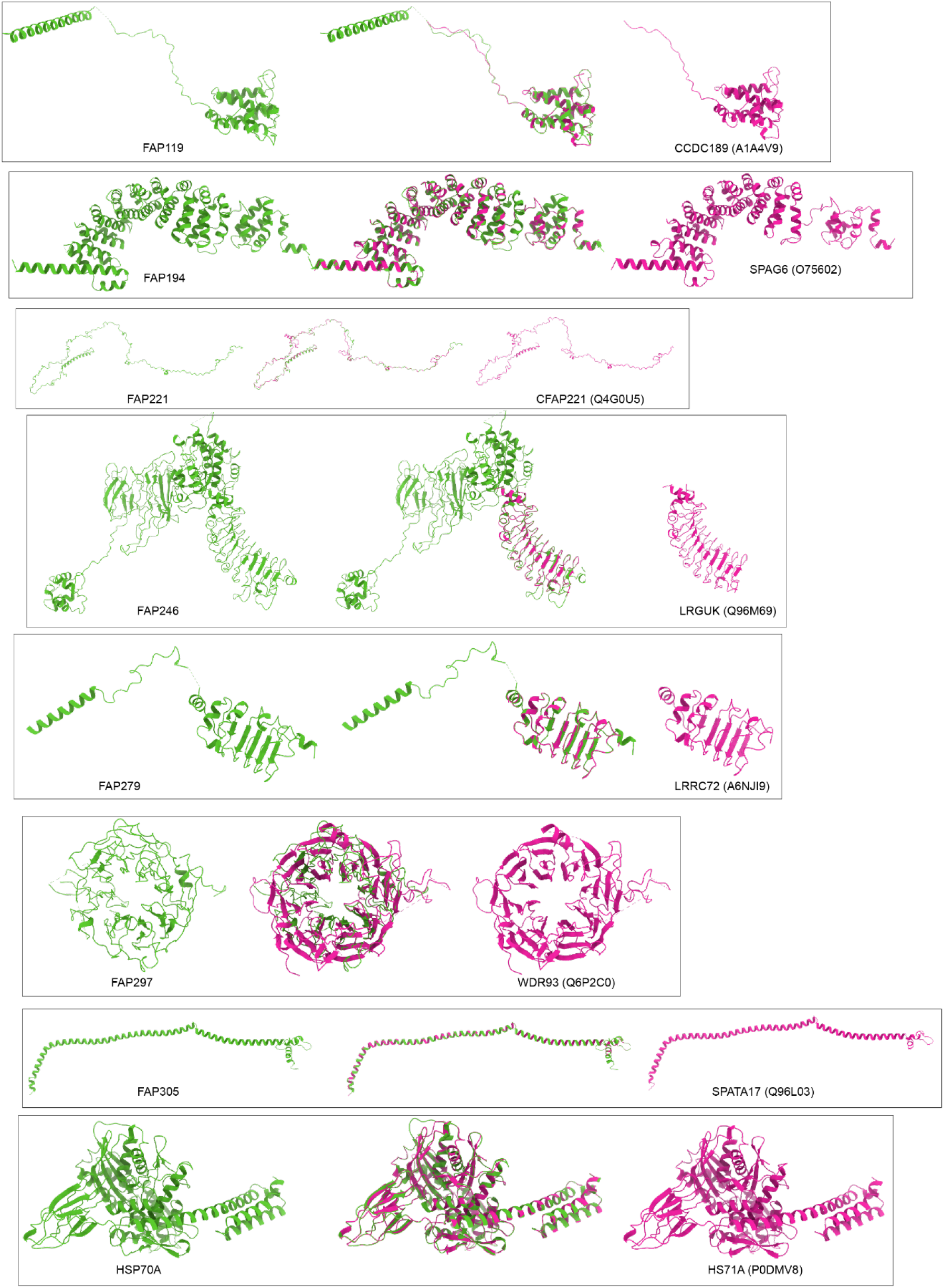
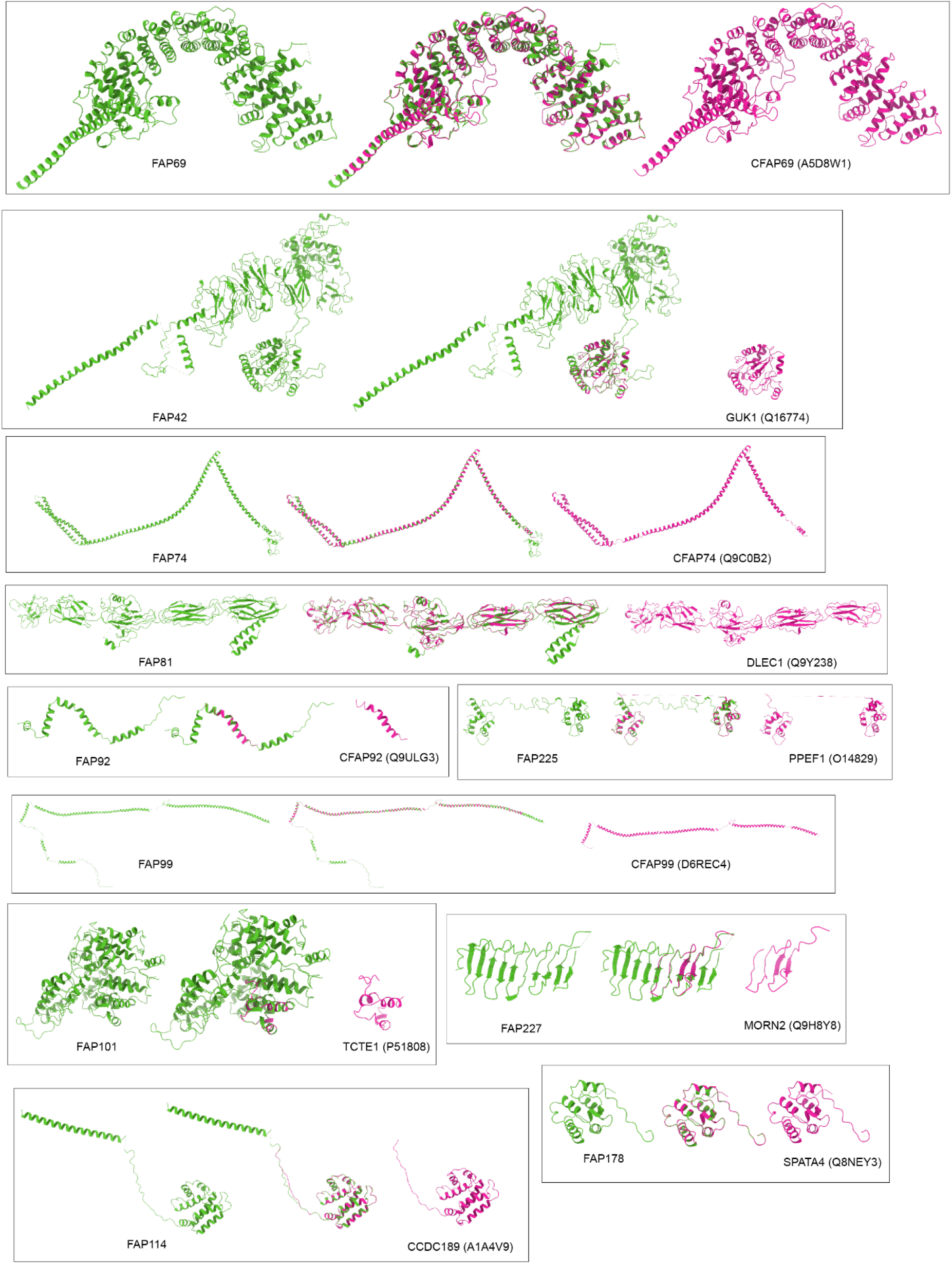
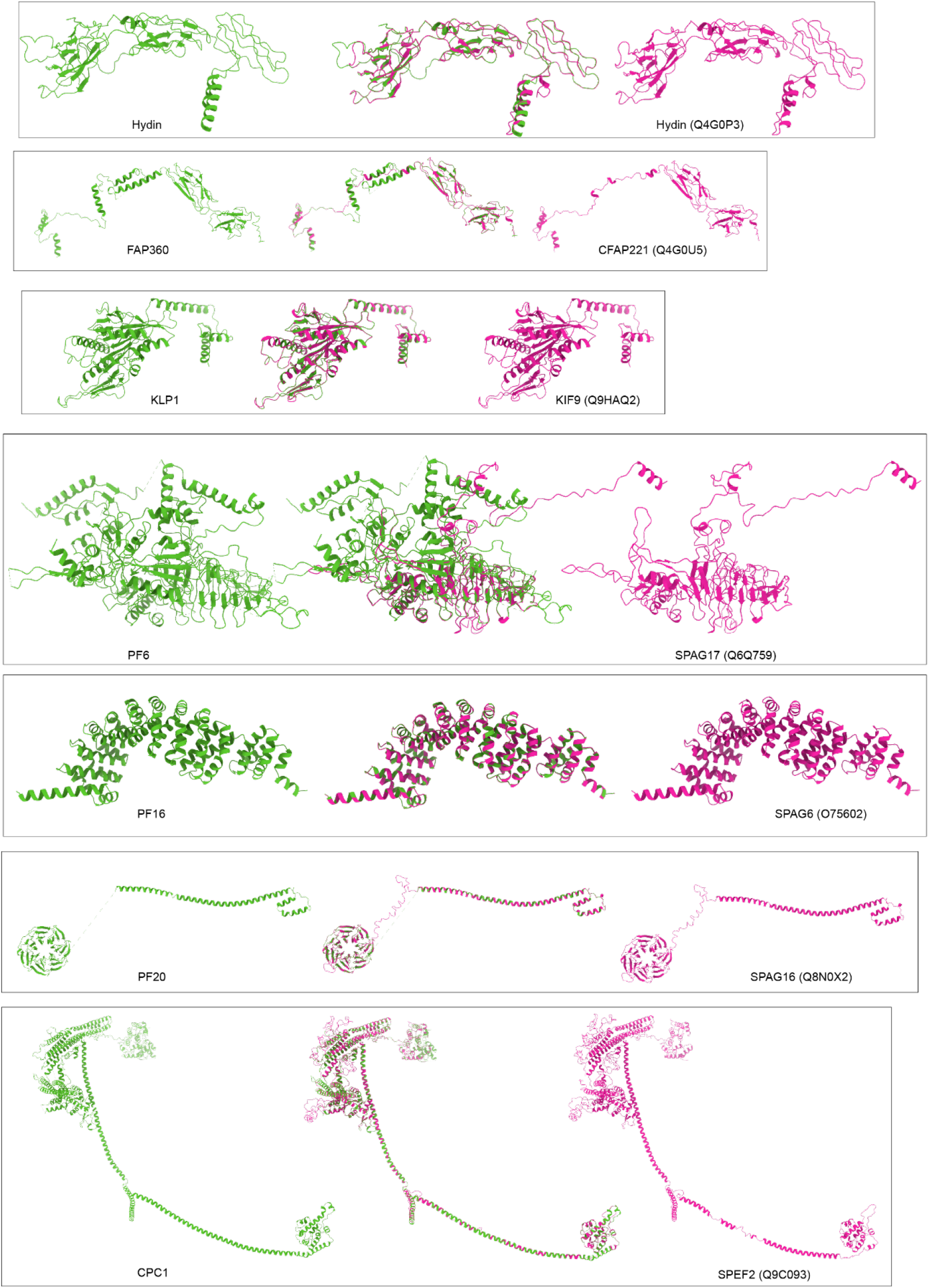

All human orthologs were searched out using blastp (https://blast.ncbi.nlm.nih.gov/Blast.cgi?PROGRAM=blastp&PAGE_TYPE=BlastSearch&LINK_LOC=blasthome). The best matched candidates were further verified by comparing with the listed human orthologs from the Chlamydomonas Flagellar Proteome Project (http://chlamyfp.org/)^1^. The atomic structures of these human orthologs were modeled using SWISS-MODEL (https://swissmodel.expasy.org/interactive#structure) using the models of *C. reinhardtii* CA determined in our study. In each box, the green molecule (left) represents the experimentally determined structure of CA protein from *C. reinhardtii*; the pink molecule (right) is the predicted structure of its human ortholog (UniProt accession numbers in the brackets); superimposition of each pair of atomic models is displayed in the center.

## Notes

### Competing Interest Statement

The authors have declared no competing interest.

## References

1. D. R. Mitchell. Evolution of cilia. Cold Spring Harb Perspect Biol, 9(1), 2017. ISSN 1943-0264 (Electronic) 1943-0264 (Linking). doi: 10.1101/cshperspect.a028290.

2. S. Khan and J. M. Scholey. Assembly, functions and evolution of archaella, flagella and cilia. Curr Biol, 28(6):R278–R292, 2018. ISSN 1879-0445 (Electronic) 0960-9822 (Linking). doi: 10.1016/j.cub.2018.01.085.

3. G. Langousis and K. L. Hill. Motility and more: the flagellum of trypanosoma brucei. Nature Reviews Microbiology, 12(7):505–518, 2014. ISSN 1740-1526. doi: 10.1038/nrmicro3274.

4. P. Satir and S. T. Christensen. Overview of structure and function of mammalian cilia. Annu Rev Physiol, 69:377–400, 2007. ISSN 0066-4278 (Print) 0066-4278 (Linking). doi: 10.1146/annurev.physiol.69.040705.141236.

5. B. A. Afzelius. Cilia-related diseases. Journal of Pathology, 204(4):470–477, 2004. ISSN 0022-3417. doi: 10.1002/path.1652.

6. B. Button, L. H. Cai, C. Ehre, M. Kesimer, D. B. Hill, J. K. Sheehan, R. C. Boucher, and M. Rubinstein. A periciliary brush promotes the lung health by separating the mucus layer from airway epithelia. Science, 337(6097):937–941, 2012. ISSN 0036-8075. doi: 10.1126/science.1223012.

7. S. Nonaka, Y. Tanaka, Y. Okada, S. Takeda, A. Harada, Y. Kanai, M. Kido, and N. Hirokawa. Randomization of left-right asymmetry due to loss of nodal cilia generating leftward flow of extraembryonic fluid in mice lacking kif3b motor protein. Cell, 95(6):829–837, 1998. ISSN 0092-8674. doi: Doi10.1016/S0092-8674(00)81705-5.

8. G. J. Vanaken, L. Bassinet, M. Boon, R. Mani, I. Honore, J. F. Papon, H. Cuppens, M. Jaspers, N. Lorent, A. Coste, E. Escudier, S. Amselem, B. Maitre, M. Legendre, and S. Christin-Maitre. Infertility in an adult cohort with primary ciliary dyskinesia: phenotype-gene association. European Respiratory Journal, 50(5), 2017. ISSN 0903-1936. doi: Artn170031410.1183/13993003. 00314-2017.

9. S. Yoshiba, H. Shiratori, I. Y. Kuo, A. Kawasumi, K. Shinohara, S. Nonaka, Y. Asai, G. Sasaki, J. A. Belo, H. Sasaki, J. Nakai, B. Dworniczak, B. E. Ehrlich, P. Pennekamp, and H. Hamada. Cilia at the node of mouse embryos sense fluid flow for left-right determination via pkd2. Science, 338(6104):226–231, 2012. ISSN 0036-8075. doi: 10.1126/science.1222538.

10. M. Fliegauf, T. Benzing, and H. Omran. Mechanisms of disease - when cilia go bad: cilia defects and ciliopathies. Nature Reviews Molecular Cell Biology, 8(11):880–893, 2007. ISSN 1471-0072. doi: 10.1038/nrm2278.

11. F. Hildebrandt, T. Benzing, and N. Katsanis. Ciliopathies. N Engl J Med, 364 (16):1533–43, 2011. ISSN 1533-4406 (Electronic) 0028-4793 (Linking). doi: 10.1056/NEJMra1010172.

12. J. F. Reiter and M. R. Leroux. Genes and molecular pathways underpinning ciliopathies. Nature Reviews Molecular Cell Biology, 18(9):533–547, 2017. ISSN 1471-0072. doi: 10.1038/nrm.2017.60.

13. K. H. Bui, T. Yagi, R. Yamamoto, R. Kamiya, and T. Ishikawa. Polarity and asymmetry in the arrangement of dynein and related structures in the chlamydomonas axoneme. Journal of Cell Biology, 198(5):913–925, 2012. ISSN 0021-9525. doi: 10.1083/jcb.201201120.

14. T. Ishikawa. Axoneme structure from motile cilia. Cold Spring Harbor Perspectives in Biology, 9(1), 2017. ISSN 1943-0264. doi: ARTNa02807610.1101/cshperspect.a028076.

15. D. Nicastro, C. Schwartz, J. Pierson, R. Gaudette, M. E. Porter, and J. R. McIntosh. The molecular architecture of axonemes revealed by cryoelectron tomography. Science, 313(5789):944–8, 2006. ISSN 1095-9203 (Electronic) 0036-8075 (Linking). doi: 10.1126/science.1128618.

16. J. F. Lin and D. Nicastro. Asymmetric distribution and spatial switching of dynein activity generates ciliary motility. Science, 360(6387), 2018. ISSN 0036-8075. doi: ARTNeaar196810.1126/science.aar1968.

17. T. J. Mitchison and H. M. Mitchison. Cell biology how cilia beat. Nature, 463 (7279):308–309, 2010. ISSN 0028-0836. doi: 10.1038/463308a.

18. R. Viswanadha, W. S. Sale, and M. E. Porter. Ciliary motility: Regulation of axonemal dynein motors. Cold Spring Harbor Perspectives in Biology, 9(8), 2017. ISSN 1943-0264. doi: ARTNa01832510.1101/cshperspect.a018325.

19. K. W. Foster, J. Vidyadharan, and A. Sangani. How cilia or eukaryotic flagella beat. Biophysical Journal, 108(2):459a–459a, 2015. ISSN 0006-3495. doi: DOI10.1016/j.bpj.2014.11.2504.

20. M. V. Sataric, T. Nemes, D. Sekulic, and J. A. Tuszynski. How signals of calcium ions initiate the beats of cilia and flagella. Biosystems, 182:42–51, 2019. ISSN 0303-2647. doi: 10.1016/j.biosystems.2019.103981.

21. P. Satir, T. Heuser, and W. S. Sale. A structural basis for how motile cilia beat. Bioscience, 64(12):1073–1083, 2014. ISSN 0006-3568. doi: 10.1093/biosci/biu180.

22. M. Wirschell, R. Yamamoto, L. Alford, A. Gokhale, A. Gaillard, and W. S. Sale. Regulation of ciliary motility: conserved protein kinases and phosphatases are targeted and anchored in the ciliary axoneme. Arch Biochem Biophys, 510(2):93–100, 2011. ISSN 1096-0384 (Electronic) 0003-9861 (Linking). doi: 10.1016/j.abb.2011.04.003.

23. R. Yokoyama, E. O’Toole, S. Ghosh, and D. R. Mitchell. Regulation of flagellar dynein activity by a central pair kinesin. Proc Natl Acad Sci U S A, 101(50): 17398–403, 2004. ISSN 0027-8424 (Print) 0027-8424 (Linking). doi: 10.1073/pnas.0406817101.

24. B. Baccetti. Evolutionary trends in sperm structure. Comp Biochem Physiol A Comp Physiol, 85(1):29–36, 1986. ISSN 0300-9629 (Print) 0300-9629 (Linking). doi: 10.1016/0300-9629(86)90457-3.

25. D. R. Mitchell. Speculations on the evolution of 9+2 organelles and the role of central pair microtubules. Biol Cell, 96(9):691–6, 2004. ISSN 0248-4900 (Print) 0248-4900 (Linking). doi: 10.1016/j.biolcel.2004.07.004.

26. S. Nonaka, Y. Tanaka, Y. Okada, S. Takeda, A. Harada, Y. Kanai, M. Kido, and N. Hirokawa. Randomization of left-right asymmetry due to loss of nodal cilia generating leftward flow of extraembryonic fluid in mice lacking kif3b motor protein. Cell, 95(6):829–37, 1998. ISSN 0092-8674 (Print) 0092-8674. doi: 10.1016/s0092-8674(00)81705-5.

27. S. Nonaka, Y. Tanaka, Y. Okada, S. Takeda, A. Harada, Y. Kanai, M. Kido, and N. Hirokawa. Randomization of left-right asymmetry due to loss of nodal cilia generating leftward flow of extraembryonic fluid in mice lacking kif3b motor protein (vol 95, pg 829, 95). Cell, 99(1), 1999. ISSN 0092-8674.

28. T. Yagi and M. Nishiyama. High hydrostatic pressure induces vigorous flagellar beating in chlamydomonas non-motile mutants lacking the central apparatus. Sci Rep, 10(1):2072, 2020. ISSN 2045-2322 (Electronic) 2045-2322 (Linking). doi: 10.1038/s41598-020-58832-8.

29. B. I. Carbajal-Gonzalez, T. Heuser, X. F. Fu, J. F. Lin, B. W. Smith, D. R. Mitchell, and D. Nicastro. Conserved structural motifs in the central pair complex of eukaryotic flagella. Cytoskeleton, 70(2):101–120, 2013. ISSN 1949-3584. doi: 10.1002/cm.21094.

30. T. D. Loreng and E. F. Smith. The central apparatus of cilia and eukaryotic flagella. Cold Spring Harbor Perspectives in Biology, 9(2), 2017. ISSN 1943-0264. doi: ARTNa02811810.1101/cshperspect.a028118.

31. M. E. Teves, D. R. Nagarkatti-Gude, Z. B. Zhang, and J. F. Strauss. Mammalian axoneme central pair complex proteins: Broader roles revealed by gene knockout phenotypes. Cytoskeleton, 73(1):3–22, 2016. ISSN 1949-3584. doi: 10.1002/cm.21271.

32. C. K. Omoto, I. R. Gibbons, R. Kamiya, C. Shingyoji, K. Takahashi, and G. B. Witman. Rotation of the central pair microtubules in eukaryotic flagella. Mol Biol Cell, 10(1):1–4, 1999. ISSN 1059-1524 (Print) 1059-1524 (Linking). doi: 10.1091/mbc.10.1.1.

33. C. G. DiPetrillo and E. F. Smith. The pcdp1 complex coordinates the activity of dynein isoforms to produce wild-type ciliary motility. Molecular Biology of the Cell, 22(23):4527–4538, 2011. ISSN 1059-1524. doi: 10.1091/mbc.E11-08-0739.

34. K. Kikushima. Central pair apparatus enhances outer-arm dynein activities through regulation of inner-arm dyneins. Cell Motility and the Cytoskeleton, 66(5):272–280, 2009. ISSN 0886-1544. doi: 10.1002/cm.20355.

35. T. Oda, H. Yanagisawa, T. Yagi, and M. Kikkawa. Mechanosignaling between central apparatus and radial spokes controls axonemal dynein activity. Journal of Cell Biology, 204(5):807–819, 2014. ISSN 0021-9525. doi: 10.1083/jcb. 201312014.

36. E. F. Smith. Regulation of flagellar dynein by the axonemal central apparatus. Cell Motility and the Cytoskeleton, 52(1):33–42, 2002. ISSN 0886-1544. doi: 10.1002/cm.10031.

37. M. J. Wargo and E. F. Smith. Asymmetry of the central apparatus defines the location of active microtubule sliding in chlamydomonas flagella. Proc Natl Acad Sci U S A, 100(1):137–42, 2003. ISSN 0027-8424 (Print) 0027-8424 (Linking). doi: 10.1073/pnas.0135800100.

38. F. D. Warner and P. Satir. The structural basis of ciliary bend formation. radial spoke positional changes accompanying microtubule sliding. J Cell Biol, 63 (1):35–63, 1974. ISSN 0021-9525 (Print) 0021-9525 (Linking). doi: 10.1083/jcb.63.1.35.

39. C. B. Lindemann and K. S. Kanous. “geometric clutch” hypothesis of axonemal function: key issues and testable predictions. Cell Motil Cytoskeleton, 31(1): 1–8, 1995. ISSN 0886-1544 (Print) 0886-1544 (Linking). doi: 10.1002/cm. 970310102.

40. C. B. Lindemann and K. A. Lesich. Flagellar and ciliary beating: the proven and the possible. Journal of Cell Science, 123(4):519–528, 2010. ISSN 0021-9533. doi: 10.1242/jcs.051326.

41. D. R. Mitchell and M. Nakatsugawa. Bend propagation drives central pair rotation in chlamydomonas reinhardtii flagella. Journal of Cell Biology, 166(5): 709–715, 2004. ISSN 0021-9525. doi: 10.1083/jcb.200406148.

42. R. Sapiro, I. Kostetskii, P. Olds-Clarke, G. L. Gerton, G. L. Radice, and J. F. Strauss. Male infertility, impaired sperm motility, and hydrocephalus in mice deficient in sperm-associated antigen 6. Molecular and Cellular Biology, 22 (17):6298–6305, 2002. ISSN 0270-7306. doi: 10.1128/Mcb.22.17.6298-6305. 2002.

43. M. E. Teves, P. R. Sears, W. Li, Z. G. Zhang, W. X. Tang, L. van Reesema, R. M. Costanzo, C. W. Davis, M. R. Knowles, J. F. Strauss, and Z. B. Zhang. Sperm-associated antigen 6 (spag6) deficiency and defects in ciliogenesis and cilia function: Polarity, density, and beat. Plos One, 9(10), 2014. ISSN 1932-6203. doi: ARTNe10727110.1371/journal.pone.0107271.

44. D. F. Zheng, Q. Wang, J. P. Wang, Z. Q. Bao, S. W. Wu, L. Ma, D. M. Chai, Z. P. Wang, and Y. S. Tao. The emerging role of sperm-associated antigen 6 gene in the microtubule function of cells and cancer. Molecular Therapy-Oncolytics, 15:101–107, 2019. ISSN 2372-7705. doi: 10.1016/j.omto.2019.08.011.

45. G. Fu, L. Zhao, E. Dymek, Y. Q. Hou, K. K. Song, N. Phan, Z. G. Shang, E. F. Smith, G. B. Witman, and D. Nicastro. Structural organization of the c1a-e-c supercomplex within the ciliary central apparatus. Journal of Cell Biology, 218 (12):4236–4251, 2019. ISSN 0021-9525. doi: 10.1083/jcb.201906006.

46. D. Dai, M. Ichikawa, K. Peri, R. Rebinsky, and K. H. Bui. Identification and mapping of central pair proteins by proteomic analysis. Biophysics and Physicobiology, 17:71–85, 2020. doi: 10.2142/biophysico.BSJ-2019048.

47. L. Zhao, Y. Q. Hou, T. Picariello, B. Craige, and G. B. Witman. Proteome of the central apparatus of a ciliary axoneme. Journal of Cell Biology, 218(6): 2051–2070, 2019. ISSN 0021-9525. doi: 10.1083/jcb.201902017.

48. M. Bernstein, P. L. Beech, S. G. Katz, and J. L. Rosenbaum. A new kinesin-like protein (klp1) localized to a single microtubule of the chlamydomonas flagellum. J Cell Biol, 125(6):1313–26, 1994. ISSN 0021-9525 (Print) 0021-9525 (Linking). doi: 10.1083/jcb.125.6.1313.

49. S K Dutcher, B Huang, and D J Luck. Genetic dissection of the central pair microtubules of the flagella of chlamydomonas reinhardtii. Journal of Cell Biology, 98(1):229–236, 1984. ISSN 0021-9525. doi: 10.1083/jcb.98.1.229.

50. B. F. Mitchell, L. B. Pedersen, M. Feely, J. L. Rosenbaum, and D. R. Mitchell. Atp production in chlamydomonas reinhardtii flagella by glycolytic enzymes. Mol Biol Cell, 16(10):4509–18, 2005. ISSN 1059-1524 (Print) 1059-1524 (Linking). doi: 10.1091/mbc.e05-04-0347.

51. E. F. Smith and P. A. Lefebvre. Pf20 gene product contains wd repeats and localizes to the intermicrotubule bridges in chlamydomonas flagella. Molecular Biology of the Cell, 8(3):455–467, 1997. ISSN 1059-1524.

52. M. J. Wargo, E. E. Dymek, and E. F. Smith. Calmodulin and pf6 are components of a complex that localizes to the c1 microtubule of the flagellar central apparatus. Journal of Cell Science, 118(20):4655–4665, 2005. ISSN 0021-9533. doi: 10.1242/jcs.02585.

53. D. R. Mitchell and W. S. Sale. Characterization of a chlamydomonas insertional mutant that disrupts flagellar central pair microtubule-associated structures. Journal of Cell Biology, 144(2):293–304, 1999. ISSN 0021-9525. doi: DOI10.1083/jcb.144.2.293.

54. G. Rupp, E. O’Toole, and M. E. Porter. The chlamydomonas pf6 locus encodes a large alanine/proline-rich polypeptide that is required for assembly of a central pair projection and regulates flagellar motility. Molecular Biology of the Cell, 12(3):739–751, 2001. ISSN 1059-1524. doi: DOI10.1091/mbc.12.3.739.

55. David Starling and John Randall. The flagella of temporary dikaryons of chlamydomonas reinhardii. Genetical Research, 18(1):107–113, 1971. ISSN 0016-6723 1469-5073. doi: 10.1017/s0016672300012465.

56. E. E. Levine M. A. Olmsted W. T. Ebersold, R. P. Levine. Linkage maps in chlamydomonas reinhardi. Genetics, 47(5):531–543, 1962.

57. H. Zhang and D. R. Mitchell. Cpc1, a chlamydomonas central pair protein with an adenylate kinase domain. J Cell Sci, 117(Pt 18):4179–88, 2004. ISSN 0021-9533 (Print) 0021-9533 (Linking). doi: 10.1242/jcs.01297.

58. D. K. Khona, V. G. Rao, M. J. Motiwalla, P. C. Varma, A. R. Kashyap, K. Das, S. M. Shirolikar, L. Borde, J. A. Dharmadhikari, A. K. Dharmadhikari, S. Mukhopadhyay, D. Mathur, and J. S. D’Souza. Anomalies in the motion dynamics of long-flagella mutants of chlamydomonas reinhardtii. J Biol Phys, 39(1):1–14, 2013. ISSN 0092-0606 (Print) 0092-0606 (Linking). doi: 10.1007/s10867-012-9282-8.

59. E. F. Smith and P. A. Lefebvre. Defining functional domains within pf16: A central apparatus component required for flagellar motility. Cell Motility and the Cytoskeleton, 46(3):157–165, 2000. ISSN 0886-1544. doi: Doi10.1002/1097-0169(200007)46:3<157::Aid-Cm1>3.0.Co;2-D.

60. X. D. Hu, R. C. Yan, X. R. Cheng, L. Z. Song, W. Zhang, K. K. Li, and S. T. Zhao. The function of sperm-associated antigen 6 in neuronal proliferation and differentiation. Journal of Molecular Histology, 47(6):531–540, 2016. ISSN 1567-2379. doi: 10.1007/s10735-016-9694-z.

61. B. Kobe and A. V. Kajava. When protein folding is simplified to protein coiling: the continuum of solenoid protein structures. Trends in Biochemical Sciences, 25(10):509–515, 2000. ISSN 0968-0004. doi: Doi10.1016/S0968-0004(00)01667-4.

62. C. G. DiPetrillo and E. F. Smith. Pcdp1 is a central apparatus protein that binds ca2+-calmodulin and regulates ciliary motility. Journal of Cell Biology, 189(3): 601–612, 2010. ISSN 0021-9525. doi: 10.1083/jcb.200912009.

63. I. Nakano and C. Shingyoji. Central-pair linked regulation of microtubule sliding by calcium in flagellar axonemes. Molecular Biology of the Cell, 12:314a–314a, 2001. ISSN 1059-1524.

64. E. F. Smith. Regulation of flagellar dynein by calcium and a role for an axonemal calmodulin and calmodulin-dependent kinase. Mol Biol Cell, 13 (9):3303–13, 2002. ISSN 1059-1524 (Print) 1059-1524 (Linking). doi: 10.1091/mbc.e02-04-0185.

65. I. Grossman-Haham, N. Coudray, Z. L. Yu, F. Wang, N. Zhang, G. Bhabha, and R. D. Vale. Structure of the radial spoke head and insights into its role in mechanoregulation of ciliary beating. Nature Structural Molecular Biology, 28 (1), 2021. ISSN 1545-9993. doi: 10.1038/s41594-020-00519-9.

66. S. Cindric, G. W. Dougherty, H. Olbrich, R. Hjeij, N. T. Loges, I. Amirav, M. C. Philipsen, J. K. Marthin, K. G. Nielsen, S. Sutharsan, J. Raidt, C. Werner, P. Pennekamp, B. Dworniczak, and H. Omran. Spef2-and hydin-mutant cilia lack the central pair-associated protein spef2, aiding primary ciliary dyskinesia diagnostics. Am J Respir Cell Mol Biol, 62(3):382–396, 2020. ISSN 1535-4989 (Electronic) 1044-1549 (Linking). doi: 10.1 165/rcmb.2019-0086OC.

67. C. F. Tu, H. C. Nie, L. L. Meng, W. L. Wang, H. Y. Li, S. M. Yuan, D. H. Cheng, W. B. He, G. Liu, J. Du, F. Gong, G. X. Lu, G. Lin, Q. J. Zhang, and Y. Q. Tan. Novel mutations in spef2 causing different defects between flagella and cilia bridge: the phenotypic link between mmaf and pcd. Human Genetics, 139(2): 257–271, 2020. ISSN 0340-6717. doi: 10.1007/s00439-020-021 10-0.

68. J. M. Brown, C. G. DiPetrillo, E. F. Smith, and G. B. Witman. A fap46 mutant provides new insights into the function and assembly of the c1d complex of the ciliary central apparatus. Journal of Cell Science, 125(16):3904–3913, 2012. ISSN 0021-9533. doi: 10.1242/jcs.107151.

69. L. Lee, D. R. Campagna, J. L. Pinkus, H. Mulhern, T. A. Wyatt, J. H. Sisson, J. A. Pavlik, G. S. Pinkus, and M. D. Fleming. Primary ciliary dyskinesia in mice lacking the novel ciliary protein pcdp1. Molecular and Cellular Biology, 28(3):949–957, 2008. ISSN 0270-7306. doi: 10.1 128/Mcb.00354-07.

70. H. Miyata, K. Shimada, A. Morohoshi, S. Oura, T. Matsumura, Z. L. Xu, Y. Oyama, and M. Ikawa. Testis-enriched kinesin kif9 is important for progressive motility in mouse spermatozoa. Faseb Journal, 34(4):5389–5400, 2020. ISSN 0892-6638. doi: 10.1096/fj.201902755R.

71. Z. Shang, K. Zhou, C. Xu, R. Csencsits, J. C. Cochran, and C. V. Sindelar. High-resolution structures of kinesin on microtubules provide a basis for nucleotide-gated force-generation. Elife, 3:e04686, 2014. ISSN 2050-084X (Electronic) 2050-084X (Linking). doi: 10.7554/eLife.04686.

72. A. Yildiz, M. Tomishige, R. D. Vale, and P. R. Selvin. Kinesin walks handover-hand. Science, 303(5658):676–8, 2004. ISSN 1095-9203 (Electronic) 0036-8075 (Linking). doi: 10.1126/science.1093753.

73. K. F. Lechtreck and G. B. Witman. Chlamydomonas reinhardtii hydin is a central pair protein required for flagellar motility. Journal of Cell Biology, 176 (4):473–482, 2007. ISSN 0021-9525. doi: 10.1083/jcb.200611115.

74. M. J. Wargo and E. F. Smith. Asymmetry of the central apparatus defines the location of active microtubule sliding in chlamydomonas flagella. Proceedings of the National Academy of Sciences of the United States of America, 100(1): 137–142, 2003. ISSN 0027-8424. doi: DOI10.1073/pnas.0135800100.

75. M. R. Leung, M. C. Roelofs, R. T. Ravi, P. Maitan, H. Henning, M. Zhang, E. G. Bromfield, S. C. Howes, B. M. Gadella, H. Bloomfield-Gadelha, and T. Zeev-Ben-Mordehai. The multi-scale architecture of mammalian sperm flagella and implications for ciliary motility. EMBO J, 40(7):e107410, 2021. ISSN 1460-2075 (Electronic) 0261-4189 (Linking). doi: 10.15252/embj.2020107410.

76. B. Craige, J. M. Brown, and G. B. Witman. Isolation of chlamydomonas flagella. Curr Protoc Cell Biol, Chapter3:Unit3 41 1–9, 2013. ISSN 1934-2616 (Electronic) 1934-2616 (Linking). doi: 10.1002/0471143030.cb0341s59.

77. David R. Mitchell and Brandon Smith. Chapter 13 - Analysis of the Central Pair Microtubule Complex in Chlamydomonas reinhardtii, volume 92, pages 197–213. Academic Press, 2009. ISBN 0091-679X. doi: https://doi.org/10.1016/S0091-679X(08)92013-6.

78. D. N. Mastronarde. Automated electron microscope tomography using robust prediction of specimen movements. Journal of Structural Biology, 152(1):36–51, 2005. ISSN 1047-8477. doi: 10.1016/j.jsb.2005.07.007.

79. S. Q. Zheng, E. Palovcak, J. P. Armache, K. A. Verba, Y. F. Cheng, and D. A. Agard. Motioncor2: anisotropic correction of beam-induced motion for improved cryo-electron microscopy. Nature Methods, 14(4):331–332, 2017. ISSN 1548-7091. doi: 10.1038/nmeth.4193.

80. K. Zhang. Gctf: Real-time ctf determination and correction. Journal of Struc-turalBiology, 193(1):1–12, 2016. ISSN 1047-8477. doi: 10.1016/j.jsb.2015.11.003.

81. J. Zivanov, T. Nakane, B. O. Forsberg, D. Kimanius, W. J. H. Hagen, E. Lindahl, and S. H. W. Scheres. New tools for automated high-resolution cryo-em structure determination in relion-3. Elife, 7, 2018. ISSN 2050-084x. doi: ARTNe4216610.7554/eLife.42166.

82. A. Punjani, J. L. Rubinstein, D. J. Fleet, and M. A. Brubaker. cryosparc: algorithms for rapid unsupervised cryo-em structure determination. Nature Methods, 14(3):290–+, 2017. ISSN 1548-7091. doi: 10.1038/Nmeth.4169.

83. J. R. Kremer, D. N. Mastronarde, and J. R. McIntosh. Computer visualization of three-dimensional image data using imod. Journal of Structural Biology, 116(1):71–76, 1996. ISSN 1047-8477. doi: DOI10.1006/jsbi.1996.0013.

84. D. N. Mastronarde and S. R. Held. Automated tilt series alignment and tomographic reconstruction in imod. Journal of Structural Biology, 197(2):102–113, 2017. ISSN 1047-8477. doi: 10.1016/j.jsb.2016.07.011.

85. E. F. Pettersen, T. D. Goddard, C. C. Huang, G. S. Couch, D. M. Greenblatt, E. C. Meng, and T. E. Ferrin. Ucsf chimera - a visualization system for exploratoryresearch and analysis. Journal of Computational Chemistry, 25(13): 1605–1612, 2004. ISSN 0192-8651. doi: 10.1002/jcc.20084.

86. Q. Rao, L. Han, Y. Wang, P. Chai, Y. W. Kuo, R. Yang, F. Hu, Y. Yang, J. Howard, and K. Zhang. Structures of outer-arm dynein array on microtubule doublet reveal a motor coordination mechanism. Nat Struct Mol Biol, 28(10):799–810, 2021. ISSN 1545-9985. doi: 10.1038/s41594-021-00656-9.

87. A. Casanal, B. Lohkamp, and P. Emsley. Current developments in coot for macromolecular model building of electron cryo-microscopy and crystallographic data. Protein Science, 29(4):1069–1078, 2020. ISSN 0961-8368. doi: 10.1002/pro.3791.

88. P. Emsley and K. Cowtan. Coot: model-building tools for molecular graphics. Acta Crystallographica Section D-Structural Biology, 60:2126–2132, 2004. ISSN 2059-7983. doi: 10.1107/S0907444904019158.

89. G. J. Pazour, N. Agrin, J. Leszyk, and G. B. Witman. Proteomic analysis of a eukaryotic cilium. Journal of Cell Biology, 170(1):103–113, 2005. ISSN 0021-9525. doi: 10.1083/jcb.200504008.

90. G. N. Murshudov, P. Skubak, A. A. Lebedev, N. S. Pannu, R. A. Steiner, R. A. Nicholls, M. D. Winn, F. Long, and A. A. Vagin. Refmac5 for the refinement of macromolecular crystal structures. Acta Crystallographica Section D-Structural Biology, 67:355–367, 2011. ISSN 2059-7983. doi: 10.1107/S0907444911001314.

91. P. V. Afonine, B. K. Poon, R. J. Read, O. V. Sobolev, T. C. Terwilliger, A. Urzhumtsev, and P. D. Adams. Real-space refinement in phenix for cryoem and crystallography. Acta Crystallogr D Struct Biol, 74(Pt 6):531–544, 2018. ISSN 2059-7983 (Electronic) 2059-7983 (Linking). doi: 10.1107/S2059798318006551.

92. T. D. Goddard, C. C. Huang, E. C. Meng, E. F. Pettersen, G. S. Couch, J. H. Morris, and T. E. Ferrin. Ucsf chimerax: Meeting modern challenges in visualization and analysis. Protein Sci, 27(1):14–25, 2018. ISSN 1469-896X (Electronic) 0961-8368 (Linking). doi: 10.1002/pro.3235.

93. J. Schindelin, I. Arganda-Carreras, E. Frise, V. Kaynig, M. Longair, T. Pietzsch, S. Preibisch, C. Rueden, S. Saalfeld, B. Schmid, J. Y. Tinevez, D. J. White, V. Hartenstein, K. Eliceiri, P. Tomancak, and A. Cardona. Fiji: an open-source platform for biological-image analysis. Nat Methods, 9(7):676–82, 2012. ISSN 1548-7105 (Electronic) 1548-7091 (Linking). doi: 10.1038/nmeth.2019.

## References

1 Pazour, G. J., Agrin, N., Leszyk, J. & Witman, G. B. Proteomic analysis of a eukaryotic cilium. Journal of Cell Biology 170, 103–113, doi:10.1083/jcb.200504008 (2005).

